# PRC1 catalytic activity is central to Polycomb system function

**DOI:** 10.1101/667667

**Authors:** Neil P. Blackledge, Nadezda A. Fursova, Jessica R. Kelley, Miles K. Huseyin, Angelika Feldmann, Robert J. Klose

**Author notes:** These authors contributed equally to this work (Neil P. Blackledge, Nadezda A. Fursova).

## Abstract

The Polycomb repressive system is an essential chromatin-based regulator of gene expression. Despite being extensively studied, how its target genes are selected and whether its histone modifying activities are required for transcriptional repression remains controversial. Here, we directly test the requirement for PRC1 catalytic activity in Polycomb system function. We demonstrate that a mutation widely used to disrupt PRC1 catalysis is hypomorphic, complicating the interpretation of previous studies. To overcome this, we develop a new inducible mutation system in embryonic stem cells that completely ablates PRC1 catalytic activity, revealing that catalysis by PRC1 drives Polycomb chromatin domain formation and higher-order chromatin interactions. In the absence of catalysis, we uncover the primary DNA-based targeting determinants that direct Polycomb target site selection. Finally, we discover that Polycomb-mediated gene repression requires PRC1 catalytic activity. Together these discoveries provide compelling new evidence supporting a PRC1-initiated pathway for Polycomb system function in gene regulation.

## Introduction

Eukaryotic DNA is wrapped around histone octamers to form nucleosomes and chromatin which organise DNA within the confines of the nucleus. In addition to this essential packaging role, histones are also post-translationally modified and this has been proposed to regulate important chromatin-based processes (Atlasi and Stunnenberg, 2017; Kouzarides, 2007; Musselman et al., 2012b). For example, removal of enzymes that modify histones around gene promoters can lead to alterations in gene expression (Bannister and Kouzarides, 2011). However, these enzymes often contain multiple highly conserved domains, some of which are not required for catalysis, and typically form large multi-protein complexes (DesJarlais and Tummino, 2016; Schuettengruber et al., 2017). This has made it challenging to understand the extent to which the catalytic activity of histone modifying enzymes and the modifications that they place contribute to nuclear processes such as transcription.

The Polycomb repressive system is an essential regulator of developmental gene expression (reviewed in Blackledge et al., 2015; Di Croce and Helin, 2013; Schuettengruber et al., 2017). Polycomb group (PcG) proteins typically belong to one of two multiprotein complexes that have chromatin modifying activity: Polycomb Repressive Complex 1 (PRC1) is an E3 ubiquitin ligase that mono-ubiquitylates histone H2A at lysine 119 (to form H2AK119ub1) (de Napoles et al., 2004; Wang et al., 2004a) and Polycomb Repressive Complex 2 (PRC2) is a methyltransferase that mono-, di- and tri-methylates histone H3 at lysine 27 (to form H3K27me1, me2 and me3) (Cao et al., 2002; Czermin et al., 2002; Kuzmichev et al., 2002; Muller et al., 2002). PRC1 and PRC2 can recognize target gene promoters and form Polycomb chromatin domains that are characterised by high-level enrichment of these complexes and the histone modifications that they deposit (Mikkelsen et al., 2007). Perturbations in Polycomb repressive complexes can lead to alterations in the levels of H2AK119ub1 and H3K27me3, and misexpression of Polycomb target genes (Boyer et al., 2006; Endoh et al., 2008; Endoh et al., 2017; Fursova et al., 2019; Leeb et al., 2010; Pasini et al., 2007; Rose et al., 2016). In turn, these molecular pathologies can cause developmental abnormalities and other human disease states (Pasini and Di Croce, 2016; Poynter and Kadoch, 2016; Richly et al., 2011).

It has been widely proposed that Polycomb chromatin domains are formed via a PRC2-initiated mechanism, in which PRC2 is actively recruited to Polycomb target genes via DNA binding accessory subunits, transcription factors or non-coding RNAs (Brockdorff, 2013; Dietrich et al., 2012; Herranz et al., 2008; Li et al., 2010; Li et al., 2017; Perino et al., 2018) and deposits H3K27me3 that is recognised by chromodomain-containing CBX proteins within so-called canonical PRC1 (cPRC1) complexes (Cao et al., 2002; Min et al., 2003; Wang et al., 2004b). cPRC1 complexes can support local chromatin compaction and long-range interactions between Polycomb chromatin domains, which has been proposed to underpin Polycomb-mediated gene repression (Eskeland et al., 2010; Francis et al., 2004; Grau et al., 2011). However, we and others have recently provided evidence for an alternative PRC1-initiated mechanism for Polycomb chromatin domain formation. This relies on a class of variant PRC1 (vPRC1) complexes which have DNA binding activities and do not require H3K27me3 for target site recognition (Blackledge et al., 2014; Blackledge et al., 2015; Cooper et al., 2014; Endoh et al., 2017; Tavares et al., 2012). In this alternative model, we propose that, in the absence of transcription, deposition of H2AK119ub1 at gene promoters by vPRC1 complexes drives PRC2 occupancy, Polycomb chromatin domain formation and protection against aberrant gene activation events.

A fundamental distinction between the PRC2 and PRC1-initiated models for Polycomb chromatin domain formation lies in their requirement for PRC1 catalysis and H2AK119ub1. In the PRC2-initiated model, PRC1 catalytic activity is proposed to be dispensable for recruitment of cPRC1 complexes, long-range chromatin interactions, and gene repression (Eskeland et al., 2010; Illingworth et al., 2015; Kundu et al., 2017). In contrast, the PRC1-initiated model relies on H2AK119ub1 as the driving force behind Polycomb chromatin domain formation and ultimately gene repression (Blackledge et al., 2014; Cooper et al., 2014). Previous attempts to test the requirement for PRC1 catalysis in Polycomb-mediated gene repression in mammals have yielded conflicting outcomes (Endoh et al., 2012; Eskeland et al., 2010; Illingworth et al., 2015; Kundu et al., 2017), and as such its contribution to Polycomb system function remains controversial. Therefore, despite the vertebrate Polycomb system being one of the most extensively studied chromatin-based gene regulation paradigms, there is still an active debate as to whether PRC1-or PRC2-initiated pathways drive Polycomb chromatin domain formation and transcriptional repression (Blackledge et al., 2015; Schuettengruber et al., 2017; Simon and Kingston, 2009; Steffen and Ringrose, 2014). As a consequence, the mechanisms that underpin Polycomb-mediated gene repression remain elusive.

Here, we systematically dissect the role of PRC1 catalysis in Polycomb-mediated gene regulation. Using reconstituted PRC1 complexes and *in vitro* ubiquitylation assays, coupled with *in vivo* experiments in mouse embryonic stem cells (ESCs), we reveal that mutations previously employed to disrupt catalysis by PRC1 are hypomorphic and do not eliminate H2AK119ub1 in cells. We develop a new catalytic mutant of PRC1 that completely ablates E3 ubiquitin ligase activity *in vitro* and use this as a basis to develop a sophisticated conditional point mutant system that rapidly converts PRC1 into a catalytically inactive form *in vivo*. This leaves PRC1 complexes intact, but completely abrogates H2AK119ub1. Using this system, we discover that catalysis by PRC1 is a central requirement for PRC2 binding and activity at Polycomb target sites, as well as occupancy of cPRC1 complexes and long-range chromatin interactions that they mediate. Furthermore, in the absence of PRC1 catalytic activity, we reveal the existence of a DNA-based sampling mechanism inherent to vPRC1 complexes that underpins Polycomb target site selection and Polycomb chromatin domain formation. Finally, and most importantly, we discover that loss of PRC1 catalysis largely phenocopies the gene expression defects caused by complete removal of PRC1. Together, these discoveries reveal a fundamental requirement for the catalytic activity of PRC1 in Polycomb chromatin domain formation and gene repression in ESCs, supporting a PRC1-initiated mechanism for Polycomb system function in gene regulation.

## Results

### Establishing a catalytically inactive PRC1 complex

To test the importance of PRC1 E3 ubiquitin ligase activity for Polycomb system function, we required a highly validated mutant version of PRC1 completely lacking catalytic activity. PRC1 complexes form around either RING1A or RING1B, which dimerise with one of six PCGF proteins (PCGF1-6) (Gao et al., 2012; Kloet et al., 2016). cPRC1 complexes utilize PCGF2 or 4, whereas vPRC1-specific complexes incorporate PCGF1/3/5/6. To achieve catalysis, RING1A/B interact with an E2 ubiquitin-conjugating enzyme to promote transfer of ubiquitin to H2AK119, with the PCGF and other PRC1 complex components regulating this activity (Buchwald et al., 2006; Rose et al., 2016). Structural information detailing how a minimal cPRC1 complex, composed of PCGF4 and RING1B, interfaces with its cognate E2 enzyme UbcH5c was originally used to define a catalytic mutation in which isoleucine 53 of RING1B was changed to alanine (I53A) (Figure 1A) (Bentley et al., 2011; Buchwald et al., 2006). This mutation disrupts the interaction between RING1B and the E2, resulting in loss of E3 ubiquitin ligase activity *in vitro*. Based on these *in vitro* observations, RING1B^I53A^ has been used in a number of *in vivo* studies to examine the role of catalysis by PRC1, with most concluding that PRC1 catalytic activity contributes modestly to Polycomb system function (Cohen et al., 2018; Endoh et al., 2012; Eskeland et al., 2010; Illingworth et al., 2015; Pengelly et al., 2015). However, we and others have recently demonstrated that vPRC1 complexes are far more catalytically active than cPRC1 complexes, both *in vitro* and *in vivo* (Fursova et al., 2019; Gao et al., 2012; Morey et al., 2013; Rose et al., 2016). Therefore, we wanted to test whether RING1B^I53A^ is indeed catalytically inactive in the context of a more enzymatically competent PRC1 complex. To this end we reconstituted the highly active PCGF1/RING1B/RYBP vPRC1 complex containing either wild-type RING1B or RING1B^I53A^ (Figure 1B). We then carried out *in vitro* ubiquitylation assays on reconstituted nucleosome substrates. As expected, wild-type PRC1 ubiquitylated histone H2A efficiently (Figure 1C and D). However, surprisingly, complexes containing RING1B^I53A^ still ubiquitylated H2A, albeit to reduced levels (Figure 1C and D). This indicates that RING1B^I53A^-PRC1, at least *in vitro*, is hypomorphic and retains some catalytic activity.

**Figure 1.**
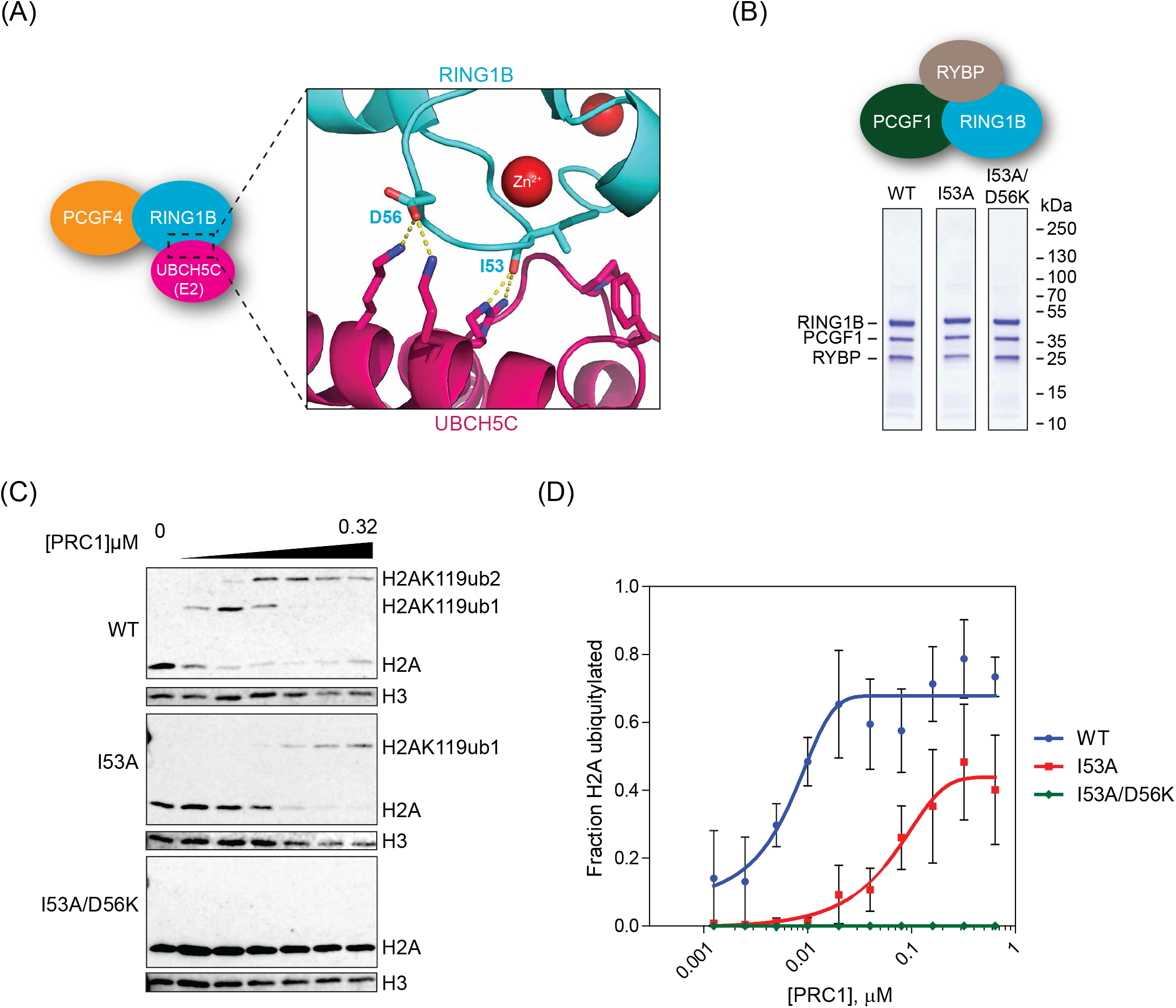
Establishing a catalytically inactive PRC1 complex *in vitro*. **(A)** A schematic of the crystal structure illustrating the RING1B-PCGF4 dimer bound to E2 ubiquitin conjugating enzyme (UBCH5C) (PDB ID: 3RPG) (Bentley et al., 2011). The key amino acids that mediate RING1B-E2 interaction are indicated. **(B)** Coomassie-stained gels of affinity-purified RING1B-PCGF1-RYBP complexes incorporating wild-type or mutated forms of RING1B. **(C)** *In vitro* E3 ubiquitin ligase assays carried out using reconstituted nucleosomes. Conversion of unmodified H2A to more slowly migrating ubiquitylated forms were measured by western blot with an H2A-specific antibody and an H3-specific antibody was used as a control. **(D)** Quantification of mean H2A ubiquitylation across a wide range of PRC1 concentrations from at least two independent experiments (C) with error bars showing SEM.

To develop a version of PRC1 which completely lacks catalytic activity, we decided to combine the I53A mutation with a second mutation, aspartate 56 to lysine (D56K), which was also shown to interfere with the RING1B-E2 interaction (Bentley et al., 2011) (Figure 1A). Importantly, in our *in vitro* E3 ubiquitin ligase assay, RING1B^I53A/D56K^-PRC1 produced no detectable H2A ubiquitylation (Figure 1C and D). Together these *in vitro* experiments demonstrate that RING1B^I53A^ is hypomorphic, whereas RING1B^I53A/D56K^ is catalytically inactive. Importantly, these observations suggest that the interpretation of experiments in which RING1B^I53A^ has been used to inactivate PRC1 require reconsideration.

### RING1B catalytic mutations are not sufficient to eliminate H2AK119ub1

Having characterised the activity of RING1B^I53A^ and RING1B^I53A/D56K^-PRC1 complexes *in vitro*, we next wanted to examine how these mutations affected H2AK119ub1 in cells. Therefore, we used CRISPR/Cas9-mediated genome engineering to introduce I53A or I53A/D56K mutations into endogenous *Ring1b* in mouse ESCs (Figure 2A). Western blot analysis revealed that RING1B levels were similar in RING1B^WT^, RING1B^I53A^ and RING1B^I53A/D56K^ cells (Figure 2B), and co-immunoprecipitation showed that PRC1 complexes with wild-type or mutated RING1B formed normally (Figure 2C). We then measured total H2AK119ub1 levels by western blot. Remarkably, this revealed that about 30% of H2AK119ub1 remained in the RING1B^I53A^ cells, with RING1B^I53A/D56K^ cells showing an additional modest, yet significant, reduction in H2AK119ub1 to about 20% of wild type levels (Figure 2B).

**Figure 2.**
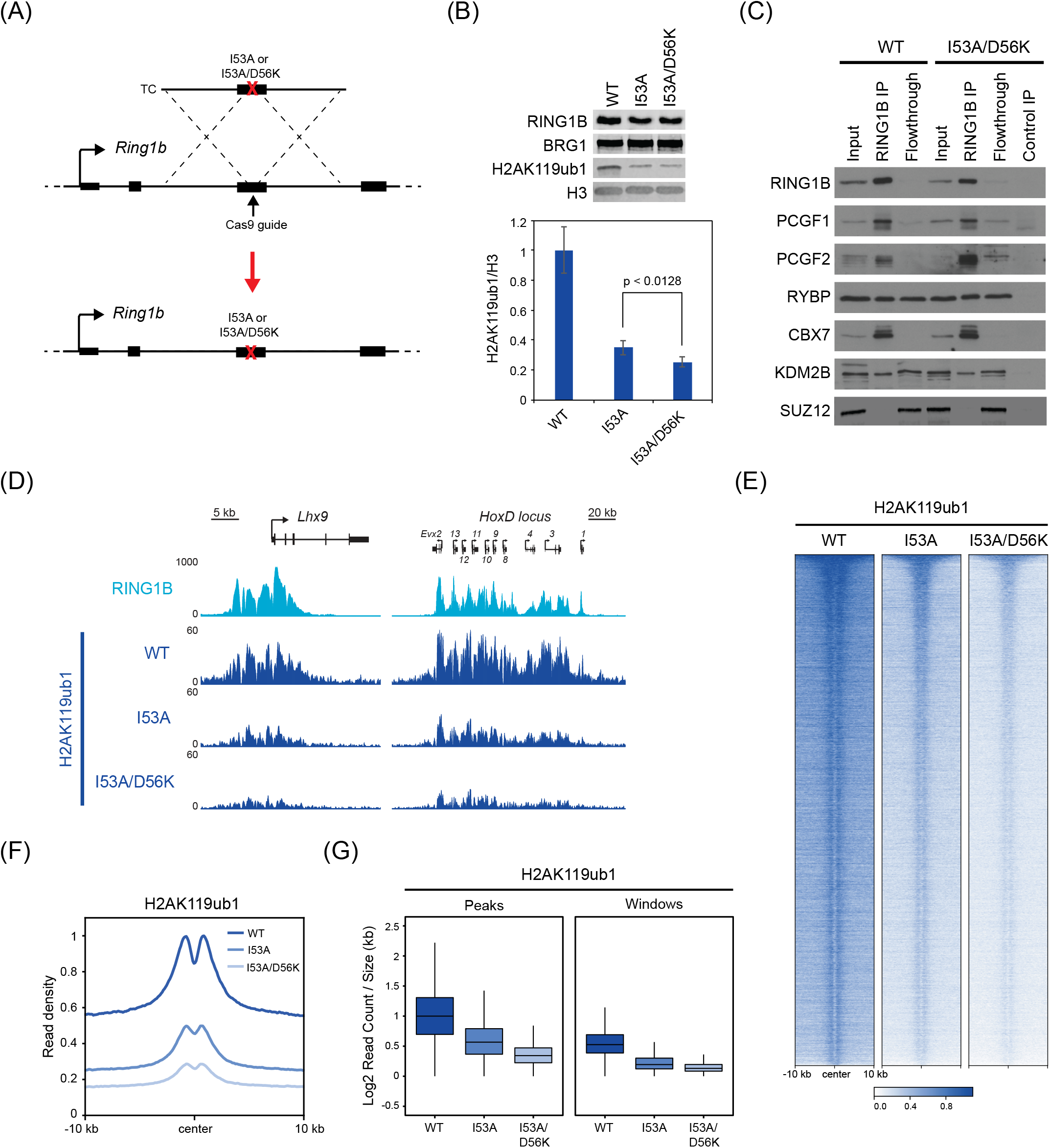
Catalytic mutations in RING1B are not sufficient to eliminate H2AK119ub1. **(A)** A schematic illustrating how I53A or I53A/D56K mutations were introduced into the endogenous *Ring1b* gene in mouse ESCs (TC = targeting construct). **(B)** Western blot analysis of RING1B (with BRG1 as a loading control) and H2AK119ub1 (with H3 as a loading control) in RING1B^WT^, RING1B^I53A^ and RING1B^I53A/D56K^ ESCs (Top panels). Quantification of H2AK119ub1 western blot data (n = 6, Bottom panel). Error bars show SEM and the P-value denotes statistical significance calculated by a paired one-tailed Student’s t-test. **(C)** Immunoprecipitation of RING1B from RING1B^WT^ or RING1B^I53A/D56K^ ESCs followed by western blot for cPRC1 and vPRC1 components as indicated. Western blot for SUZ12 (a PRC2 component) was used as a negative control. For RING1B^I53A/D56K^ ESCs, a control IP was performed with an isotype control antibody. **(D)** Genomic snapshots of classical RING1B-occupied loci, showing cChIP-seq data for RING1B in wild-type cells, and cChIP-seq for H2AK119ub1 in RING1B^WT^, RING1B^I53A^, and RING1B^I53A/D56K^ ESCs. **(E)** Heatmap analysis of H2AK119ub1 cChIP-seq at RING1B-bound sites in RING1B^WT^, RING1B^I53A^, and RING1B^I53A/D56K^ ESCs. Genomic regions were sorted based on RING1B occupancy in untreated PRC1^CKO^ ESCs. **(F)** Metaplot analysis of H2AK119ub1 cChIP-seq at RING1B-bound sites in RING1B^WT^, RING1B^I53A^, and RING1B^I53A/D56K^ ESCs. **(G)** Box plots comparing the normalised H2AK119ub1 cChIP-seq signal at RING1B-bound sites and in 100 kb windows covering the genome in RING1B^WT^, RING1B^I53A^, and RING1B^I53A/D56K^ ESCs.

We next used calibrated chromatin immunoprecipitation sequencing (cChIP-seq) to profile H2AK119ub1 in RING1B^WT^, RING1B^I53A^, and RING1B^I53A/D56K^ cells (Bonhoure et al., 2014; Hu et al., 2015; Orlando et al., 2014). In agreement with the western blot analysis, this revealed that levels of H2AK119ub1 were majorly reduced in RING1B^I53A^ cells and further reduced in RING1B^I53A/D56K^ cells, but it was not completely lost (Figure 2D-G). This was evident both for RING1B-bound sites with high levels of H2AK119ub1 (Figure 2E and F) and the low-level blanket of H2AK119ub1 that we recently demonstrated covers the genome (Figure 2G) (Fursova et al., 2019). Importantly, in both RING1B^I53A^ and RING1B^I53A/D56K^ ESCs, RING1B target sites retained considerable levels of H2AK119ub1 (Figure 2E and F). Given that RING1B^I53A/D56K^-PRC1 showed no apparent enzymatic activity *in vitro*, we were surprised to observe that such high levels of H2AK119ub1 were retained in RING1B^I53A/D56K^ cells. We hypothesised that this remaining H2AK119ub1 may come from the activity of the RING1B paralogue, RING1A, which is also expressed in ESCs, albeit at lower levels (Endoh et al., 2008). To test this possibility, we set out to introduce analogous catalytically inactivating mutations into RING1A (I50A/D53K). Generation of RING1A^I50A/D53K^ ESCs was highly efficient, but when we tried to combine RING1A^I50A/D53K^ with RING1B^I53A/D56K^, no RING1A^I50A/D53K^;RING1B^I53A/D56K^ ESCs were obtained.

In summary, our western blot and cChIP-seq analyses demonstrate that a considerable amount of H2AK119ub1 is retained in RING1B^I53A^ ESCs, indicating that this mutation cannot be used to draw conclusions on the requirement for PRC1 catalysis and H2AK119ub1 in Polycomb system function. Furthermore, the retention of H2AK119ub1 in the RING1B^I53A/D56K^ line indicates that RING1B-containing PRC1 complexes do not account for all H2AK119ub1 in ESCs and that RING1A must contribute significantly to the catalytic activity of PRC1. Our inability to isolate cells with both RING1A^I50A/D53K^ and RING1B^I53A/D56K^ suggests that PRC1 catalysis may be essential for ESC viability and Polycomb system function.

### A conditional point mutant system to inactivate PRC1 catalysis

Since we were unable to generate constitutive RING1A^I50A/D53K^;RING1B^I53A/D56K^ mutant cells, we needed to develop a new system to study whether PRC1 catalysis is required for Polycomb system function *in vivo*. We reasoned that this could be achieved by generating a cell line in which removal of catalysis could be conditionally induced. To do this, we engineered both endogenous *Ring1b* alleles to contain an I53A/D56K mutant version of the exon encoding the E2 interaction domain in the antisense orientation downstream of the corresponding wild-type exon (Figure 3A and Figure S1C). This wild-type/mutant exon pair was flanked by inverted double LoxP/Lox2272 sites and the cells were engineered to express tamoxifen-inducible CRE recombinase. In the absence of tamoxifen (OHT), wild-type RING1B would be expressed, but addition of tamoxifen would lead to an inversion event causing the cells to express catalytically inactive RING1B^I53A/D56K^ (Figure 3A-B). Importantly, to eliminate the contribution of RING1A, we also constitutively inactivated both alleles of the endogenous *Ring1a* gene by introducing RING1A^I50A/D53K^ mutations (Figure 3B).

**Figure 3.**
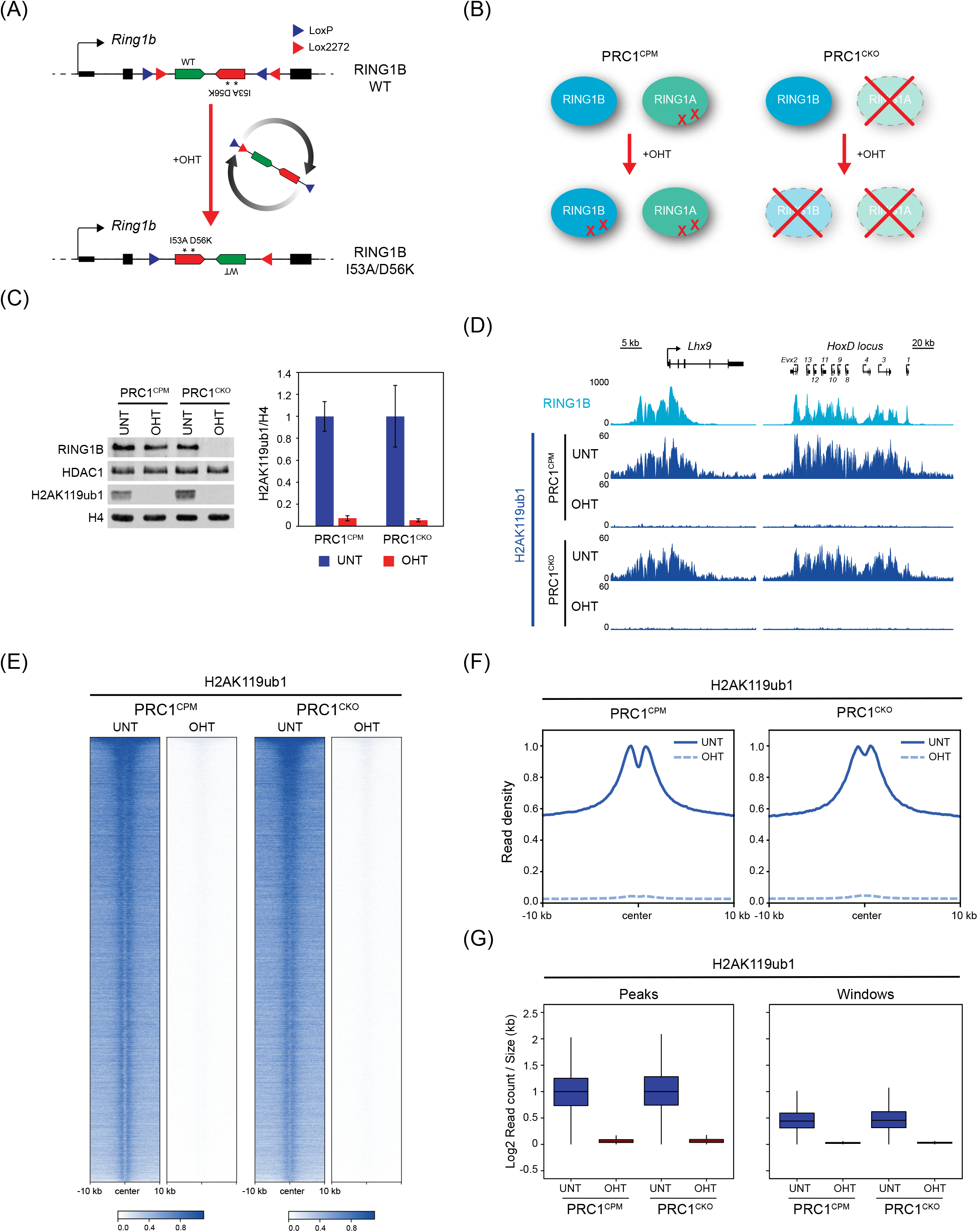
A conditional point mutant system to inactivate PRC1 catalysis. **(A)** A schematic of the engineered endogenous *Ring1b* locus in the PRC1^CPM^ system before and after OHT addition. **(B)** A schematic of the PRC1^CPM^ and PRC1^CKO^ cell lines. In PRC1^CPM^ cells, RING1A is constitutively mutated (RING1A^I50A/D53K^), and OHT addition causes conversion of RING1B into catalytically inactive form (RING1B^I53A/D56K^). In PRC1^CKO^ cells, RING1A is constitutively deleted and OHT addition causes deletion of RING1B. **(C)** Western blot analysis of RING1B (with HDAC1 as a loading control) and H2AK119ub1 (with H4 as a loading control) in untreated and OHT-treated PRC1^CPM^ and PRC1^CKO^ cells (*left panel*). A quantification of H2AK119ub1 levels relative to histone H4 with error bars representing SEM (n=4) (*right panel*). **(D)** Genomic snapshots of classical RING1B-occupied loci, showing cChIP-seq data for RING1B in untreated PRC1^CKO^ cells, and cChIP-seq for H2AK119ub1 in PRC1^CPM^ and PRC1^CKO^ cells (untreated and OHT-treated). **(E)** Heatmap analysis of H2AK119ub1 cChIP-seq at RING1B-bound sites in PRC1^CPM^ and PRC1^CKO^ cells (untreated and OHT-treated). The genomic regions were sorted based on RING1B occupancy in untreated PRC1^CKO^ ESCs. **(F)** Metaplot analysis of H2AK119ub1 cChIP-seq at RING1B-bound sites in PRC1^CPM^ and PRC1^CKO^cells (untreated and OHT-treated). **(G)** Box plots comparing the normalised H2AK119ub1 cChIP-seq signal at RING1B-bound sites and in 100 kb windows covering the genome in PRC1^CPM^ and PRC1^CKO^ cells (untreated and OHT-treated). See also Figure S1.

To validate the functionality of the PRC1 conditional point mutant (PRC1^CPM^) system, we confirmed that exon inversion occurred efficiently and *Ring1b* mRNA encoding the I53A/D56K mutation was exclusively expressed after 72 hours of tamoxifen treatment (Figure S1A-C). Importantly, RING1B protein levels were largely unchanged following addition of tamoxifen, but in contrast to constitutive RING1B^I53A^ and RING1B^I53A/D56K^ mutant cells, we now observed a complete loss of H2AK119ub1 as measured by bulk histone western blot in tamoxifen-treated PRC1^CPM^ cells (Figure 3C). To ensure that H2AK119ub1 was completely abrogated genome-wide, we carried out cChIP-seq for H2AK119ub1. This demonstrated that H2AK119ub1 was no longer found at sites enriched for RING1B or throughout the genome (Figure 3D-G, Figure S1D).

In order to compare the molecular defects that arise from catalytic inactivation of PRC1 with those that manifest following complete removal of PRC1, we also developed an isogenic ESC line, in which both copies of *Ring1a* were constitutively deleted, the first coding exon of both *Ring1b* alleles was flanked by parallel LoxP sites, and the cells were engineered to express tamoxifen-inducible CRE recombinase (Figure 3B and Figure S1E). In this PRC1 conditional knockout (PRC1^CKO^) cell line, RING1B and H2AK119ub1 were completely lost following 72 hours of tamoxifen treatment (Figure 3C). Importantly, the loss of H2AK119ub1 in the tamoxifen-treated PRC1^CKO^ was identical to that observed in the PRC1^CPM^ cells, when analysed by western blot and cChIP-seq (Figure 3C and Figure 3D-G, Figure S1D and F). The combination of isogenic conditional catalytic point mutant (PRC1^CPM^) and conditional knockout (PRC1^CKO^) ESC lines (Figure 3B) now provided us with the opportunity to directly test the PRC1-initiated model for Polycomb chromatin domain formation and ultimately define whether catalysis by PRC1 is required for Polycomb system function and gene repression.

### PRC2 binding and deposition of H3K27me3 at Polycomb target sites rely on PRC1 catalytic activity

We have previously demonstrated that removal of PRC1 results in a dramatic reduction of PRC2 and H3K27me3 levels at Polycomb target genes in line with a PRC1-initiated mechanism for Polycomb chromatin domain formation (Blackledge et al., 2014; Fursova et al., 2019; Rose et al., 2016). However, much milder effects on PRC2 and H3K27me3 were observed in RING1B^I53A^ ESCs (Illingworth et al., 2015), suggesting that the catalytic activity of PRC1 may not solely dictate Polycomb chromatin domain formation. With the knowledge that RING1B^I53A^ cells retain significant amounts of H2AK119ub1, we set out to use our new PRC1^CPM^ ESCs, in which complete loss of H2AK119ub1 could be induced, to directly test whether PRC1 catalytic activity is required for normal PRC2 targeting and H3K27me3 deposition. To do this, we performed cChIP-seq for the core PRC2 component SUZ12, and H3K27me3, in the PRC1^CPM^ and PRC1^CKO^ ESCs before and after tamoxifen treatment. In untreated cells, SUZ12 binding occurred at sites also occupied by RING1B (Figure 4A, Figure S2A-B), but strikingly in the tamoxifen-treated PRC1^CPM^ cells we observed a dramatic reduction in the levels of both SUZ12 and H3K27me3 at these sites (Figure 4A, B, D, and E, Figure S2C-D). Importantly, these changes recapitulated the effects on PRC2 observed when PRC1 was removed in the PRC1^CKO^ cells (Figure 4F-G, Figure S2C-D).

**Figure 4.**
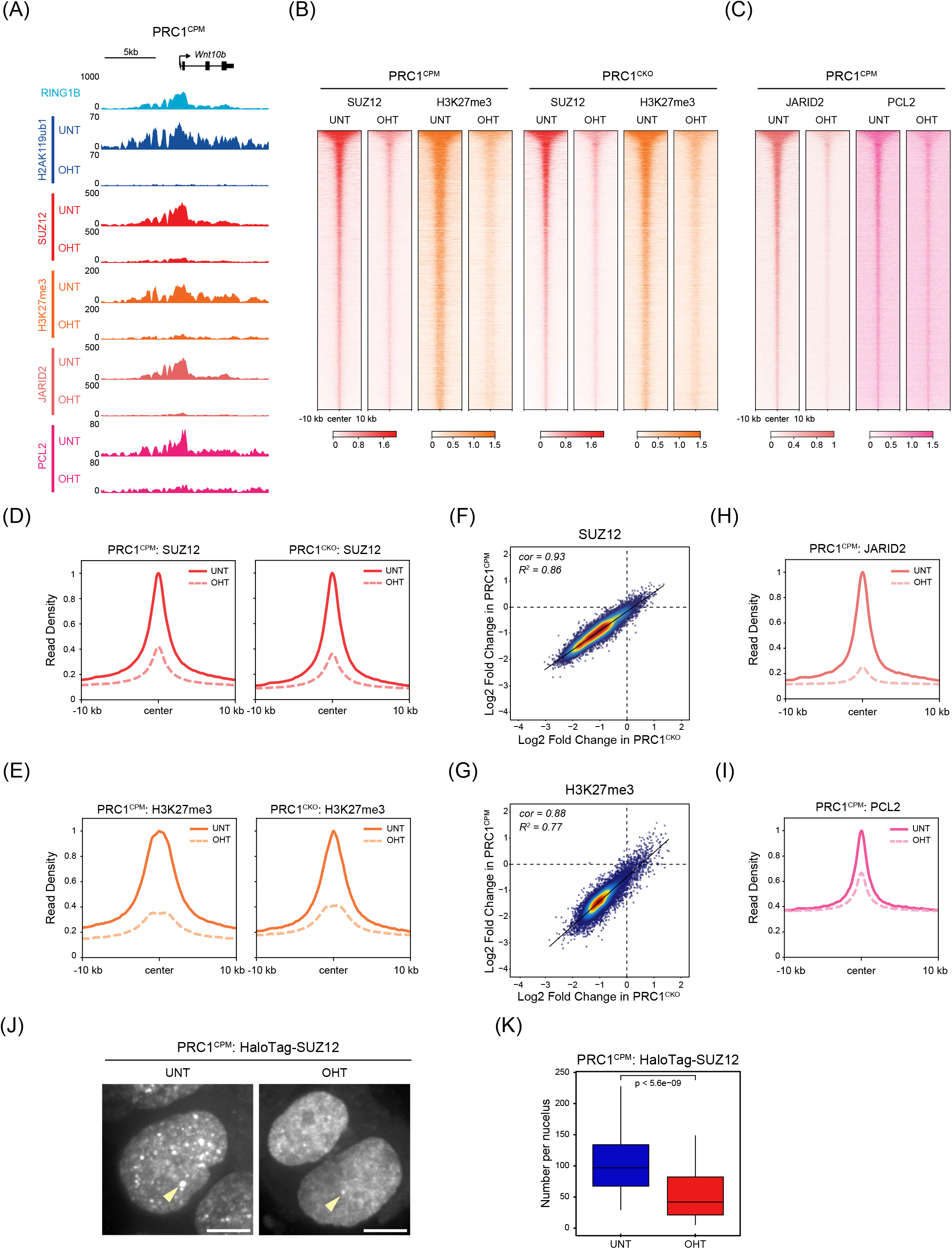
PRC2 binding and H3K27me3 deposition at Polycomb target sites rely on PRC1 catalytic activity. **(A)** Genomic snapshot of a Polycomb target gene showing cChIP-seq for H2AK119ub1, SUZ12, H3K27me3, JARID2 and PCL2 in untreated and OHT-treated PRC1^CPM^ cells. RING1B cChIP-seq in untreated cells is also shown. **(B)** Heatmap analyses of cChIP-seq for SUZ12 and H3K27me3 at PcG-occupied sites in PRC1^CPM^ and PRC1^CKO^ cells (untreated and OHT-treated). Genomic regions were sorted based on RING1B occupancy in untreated PRC1^CKO^ ESCs. **(C)** Heatmap analyses of cChIP-seq for JARID2 and PCL2 at PcG-occupied sites in PRC1^CPM^ ESCs (untreated and OHT-treated). Genomic regions were sorted as in (B). **(D)** Metaplot analysis of SUZ12 cChIP-seq at PcG-occupied sites in PRC1^CPM^ and PRC1^CKO^ cells (untreated and OHT-treated). **(E)** Metaplot analysis of H3K27me3 cChIP-seq at PcG-occupied sites in PRC1^CPM^ and PRC1^CKO^ cells (untreated and OHT-treated). **(F)** A scatter plot comparing the log2 fold changes in SUZ12 levels at PcG-occupied sites in PRC1^CPM^ versus PRC1^CKO^ ESCs following tamoxifen treatment. *R^2^* represents coefficient of determination for linear regression, and *cor* denotes Pearson correlation coefficient. **(G)** A scatter plot comparing the log2 fold changes in H3K27me3 levels at PcG-occupied sites in PRC1^CPM^ versus PRC1^CKO^ ESCs following tamoxifen treatment. *R^2^* represents coefficient of determination for linear regression, and *cor* denotes Pearson correlation coefficient. **(H)** Metaplot analysis of JARID2 cChIP-seq at PcG-occupied sites in PRC1^CPM^ cells (untreated and OHT-treated). **(I)** Metaplot analysis of PCL2 cChIP-seq at PcG-occupied sites in PRC1^CPM^ cells (untreated and OHT-treated). **(J)** Maximum intensity projections of JF_549_-Halo-SUZ12 signal in PRC1^CPM^ ESCs (untreated and OHT-treated). Examples of SUZ12 nuclear foci (Polycomb bodies) are indicated by arrowheads. Scale bar is 5 µm. **(K)** Box plots comparing the number of JF_549_-Halo-SUZ12 nuclear foci detected in PRC1^CPM^ ESCs before (n_cells_ = 55) and after OHT treatment (n_cells_ = 52). P-values denote statistical significance calculated by a Student’s t-test. See also Figure S2.

Biochemical studies have identified two subtypes of PRC2 complexes with distinct targeting modules (reviewed in Laugesen et al., 2019; Yu et al., 2019). The PRC2.1 complex contains PCL proteins, which have been shown to bind to non-methylated CpG-rich DNA and methylated histone tails (Ballare et al., 2012; Brien et al., 2012; Cai et al., 2013; Li et al., 2017; Musselman et al., 2012a; Perino et al., 2018), while the PRC2.2 complex contains the JARID2 subunit, which has been recently implicated in direct recognition of H2AK119ub1 (Cooper et al., 2016; Kalb et al., 2014). Based on the major reductions in SUZ12 binding and H3K27me3 levels in the tamoxifen-treated PRC1^CPM^ cells, we wanted to test whether occupancy of PRC2.1 and PRC2.2 was differentially affected by loss of PRC1 catalytic activity. Consistent with the proposed role of JARID2 as a reader of H2AK119ub1, cChIP-seq in PRC1^CPM^ cells revealed an almost complete loss of JARID2 binding at PcG-occupied sites following tamoxifen treatment (Figure 4A, C and H, Figure S2E). Interestingly, we also observed a reduction in JARID2 protein levels following loss of PRC1 catalytic activity (Figure S2F), suggesting that chromatin binding is required for its stability. Next, we examined the contribution of PRC1 catalytic activity to PRC2.1 occupancy by performing cChIP-seq for PCL2, a subunit specific to this complex. This revealed that in the tamoxifen-treated PRC1^CPM^ cells PCL2 occupancy at PcG-occupied sites was reduced, albeit to a lesser extent than JARID2 (Figure 4A, C and I, Figure S2E and G), while PCL2 protein levels were largely unaffected (Figure S2F). A possible explanation for this somewhat surprising result could be that, while PCL proteins confer DNA binding activity to the PRC2.1 complex, occupancy of PRC2.1 also relies on EED-dependent recognition of H3K27me3 (Margueron et al., 2009), the levels of which are strongly diminished in the absence of PRC1 catalytic activity (Figure 4A, B and E). Together, these observations demonstrate that both PRC2.1 and PRC2.2 rely heavily on PRC1 catalytic activity for binding to Polycomb target sites.

In the nucleus, PcG proteins have been shown to form cytological foci, called Polycomb bodies, that contain Polycomb-repressed genes and are highly enriched for Polycomb-associated factors (Cheutin and Cavalli, 2012; Ren et al., 2008; Saurin et al., 1998). To examine whether the major reductions in PRC2 enrichment observed at target sites in fixed cells by cChIP-seq were also evident in live cells, we added a HaloTag to the endogenous SUZ12 protein in the PRC1^CPM^ cells and used live-cell imaging to examine SUZ12 localisation in the nucleus before and after tamoxifen treatment. In untreated PRC1^CPM^ cells, we observed approximately one hundred SUZ12 nuclear foci per cell with a wide range of sizes and intensities (Figure 4J and Figure S2H). Following tamoxifen treatment and loss of PRC1 catalytic activity, the number, size, and intensity of SUZ12 foci was dramatically reduced, indicating that normal PRC2 localisation was also disrupted in live cells (Figure 4J and K, Figure S2H and I). Together, these results demonstrate that high-level PRC2 occupancy and H3K27me3 deposition at Polycomb target sites relies heavily on the catalytic activity of PRC1, supporting a PRC1-initiated mechanism for Polycomb chromatin domain formation.

### PRC1 catalytic activity drives canonical PRC1 occupancy and higher-order chromatin interactions

At Polycomb target sites, PRC2-deposited H3K27me3 is recognised by cPRC1 complexes which are proposed to mediate long-range interactions between Polycomb chromatin domains (Kundu et al., 2017; Schoenfelder et al., 2015). Previous work has reported that these long-range chromatin interactions are diminished in RING1B knockout cells but intact in RING1B^I53A^ cells (Eskeland et al., 2010; Kundu et al., 2017), leading to the conclusion that Polycomb chromatin domain interactions do not rely on catalysis by PRC1. Given the hypomorphic nature of RING1B^I53A^ cells and our new observations that H3K27me3 levels are dramatically reduced in the PRC1^CPM^ cells following tamoxifen treatment, we wanted to determine whether cPRC1 binding and higher-order chromatin interactions were affected when PRC1 catalysis was completely removed. We therefore carried out cChIP-seq for PCGF2, a core component of cPRC1 complexes in ESCs (Morey et al., 2015). In untreated PRC1^CPM^ cells, PCGF2 was enriched at RING1B-bound regions with high levels of PRC2 and H3K27me3 (Figure 5A, Figure S3A and B). Importantly, PCGF2 binding at target sites was largely lost in the tamoxifen-treated PRC1^CPM^ cells (Figure 5A-C, Figure S3C). Furthermore, protein levels of cPRC1-specific subunits were reduced following tamoxifen treatment, suggesting that a failure to bind chromatin destabilizes the cPRC1 complex (Figure 5D).

**Figure 5.**
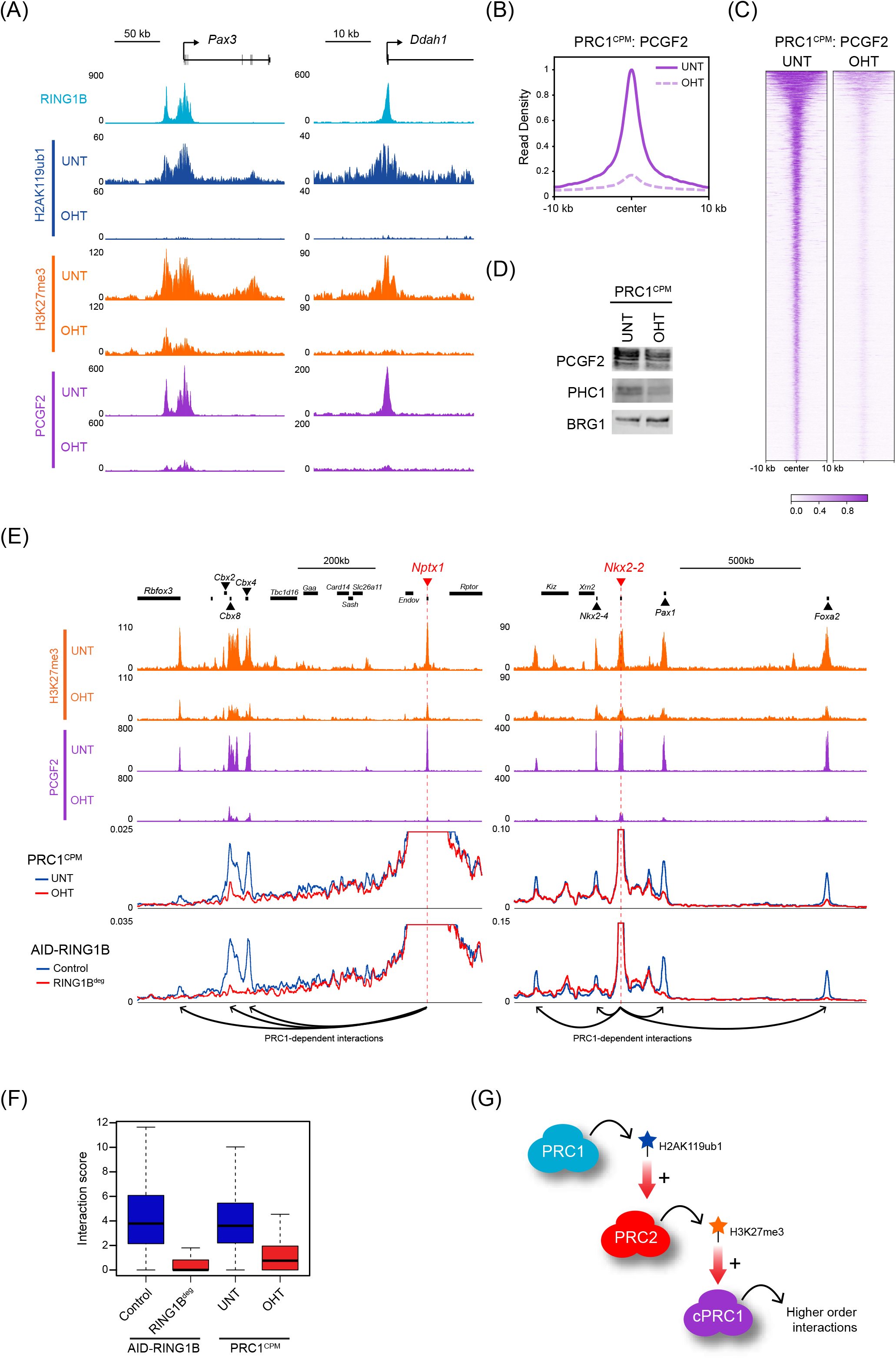
PRC1 catalytic activity drives canonical PRC1 occupancy and higher-order chromatin interactions. **(A)** Genomic snapshots of Polycomb target genes, showing cChIP-seq for H2AK119ub1, H3K27me3 and PCGF2 (cPRC1) in PRC1^CPM^ cells (untreated and OHT-treated). RING1B cChIP-seq in untreated cells is also shown. **(B)** Metaplot analysis of PCGF2 cChIP-seq at PCGF2 target sites in PRC1^CPM^ cells (untreated and OHT-treated). **(C)** Heatmap analysis of PCGF2 cChIP-seq at PCGF2 target sites in PRC1^CPM^ cells (untreated and OHT-treated). Genomic regions were sorted based on RING1B occupancy in untreated PRC1^CKO^ ESCs. **(D)** Western blot analysis of cPRC1 components (PCGF2 and PHC1) in PRC1^CPM^ cells (untreated and OHT-treated). BRG1 is shown as a loading control. **(E)** Genomic snapshots of two regions around classical Polycomb target sites used as baits in CaptureC analysis (bait capture probe positions are marked by dashed red lines). H3K27me3 and PCGF2 cChIP-seq data in PRC1^CPM^ cells (untreated and OHT-treated) are shown at the top. Below is the bait interaction landscape measured by CaptureC in PRC1^CPM^ cells (untreated and OHT-treated) and auxin-treated *Ring1a^-/-^*;AID-RING1B (RING1B^deg^) ESCs (relative to the Control cell line with intact PRC1). Long-range PRC1-dependent interactions are illustrated by arrows. **(F)** Box plot analysis of CHiCAGO scores for the interactions between the bait Polycomb target regions and other PCGF2 target sites (n = 130) in auxin-treated *Ring1a^-/-^*;AID-RING1B (RING1B^deg^) ESCs (relative to the Control cell line with intact PRC1) and PRC1^CPM^ cells (untreated and OHT-treated). **(G)** A schematic summarising the PRC1-initiated model, in which deposition of H2AK119ub1 by PRC1 drives PRC2 recruitment and H3K27me3 accumulation at Polycomb target sites, which then promotes canonical PRC1 binding and higher-order chromatin interactions between these regions. See also Figure S3.

Next, we examined whether loss of cPRC1 binding following removal of PRC1 catalytic activity had an effect on long-range interactions between Polycomb chromatin domains. To achieve this, we used a chromosome conformation capture-based approach, called CaptureC (Hughes et al., 2014), to examine interaction profiles for 24 classical Polycomb target sites that are highly enriched with PCGF2 and RING1B (Figure S3D). Strikingly, this revealed that long-range interactions formed between these regions and other classical Polycomb target sites were largely abolished in the tamoxifen-treated PRC1^CPM^ cells and this effect was highly comparable to the complete removal of PRC1 (Figure 5E and F, Figure S3E). Together, these observations further support a PRC1-initiated hierarchical recruitment model in which PRC1 catalytic activity is a primary requirement for a sequence of downstream events which culminate in the formation of Polycomb chromatin domains that can engage in long-range interactions (Figure 5G).

### DNA-based sampling activities of vPRC1 complexes drive target site selection by PRC1

The mechanisms that are responsible for the selection of target sites at which repressive Polycomb chromatin domains form remain poorly understood. cPRC1 complexes account for a significant proportion of RING1B occupancy at established Polycomb chromatin domains (Fursova et al., 2019; Morey et al., 2015), but this is a downstream consequence of PRC1 catalysis (Figure 5). Therefore, we reasoned that in the absence of PRC1 catalytic activity and cPRC1 binding, the primary mechanisms responsible for target site selection may be revealed. To examine this possibility, we carried out cChIP-seq for RING1B in the PRC1^CPM^ cells before and after tamoxifen treatment and compared RING1B occupancy to the binding of other Polycomb factors and chromatin features. In untreated PRC1^CPM^ cells, binding of RING1B correlated strongly with the levels of PCGF2 and PRC2 (SUZ12 and H3K27me3), consistent with a prominent role for cPRC1 in shaping RING1B occupancy in wild-type cells (Figure 6A, D and E and Figure S4A, E and F). Following tamoxifen treatment, RING1B binding was majorly reduced at sites that normally have high levels of PRC1 and PRC2 in untreated cells, and now correlated only modestly with cPRC1 or PRC2 occupancy (Figure 6A, D and E, Figure S4A-B and S4E-F). Surprisingly, however, RING1B levels were unchanged or even increased in tamoxifen-treated PRC1^CPM^ ESCs at a large number of sites that normally have low to moderate enrichment of PRC1 and PRC2 (Figure 6A and D and Figure S4A and B). Importantly, similar effects were observed using live-cell imaging of Polycomb foci in untreated and tamoxifen-treated PRC1^CPM^ cells in which we added a HaloTag to the endogenous RING1B protein. Following loss of PRC1 catalysis, the total number, average size and intensity of foci were only modestly reduced (Figure 6B and C, Figure S4C). However, we observed a dramatic reduction in the number of bright foci, while the number of less intense foci was largely unaffected, in agreement with our cChIP-seq analysis (Figure 6B and C, Figure S4C and D). Together this reveals that while high-level enrichment of RING1B at target sites forming Polycomb chromatin domains requires PRC1 catalytic activity, there exists low-level RING1B binding across all target sites that is independent of PRC1 catalysis.

**Figure 6.**
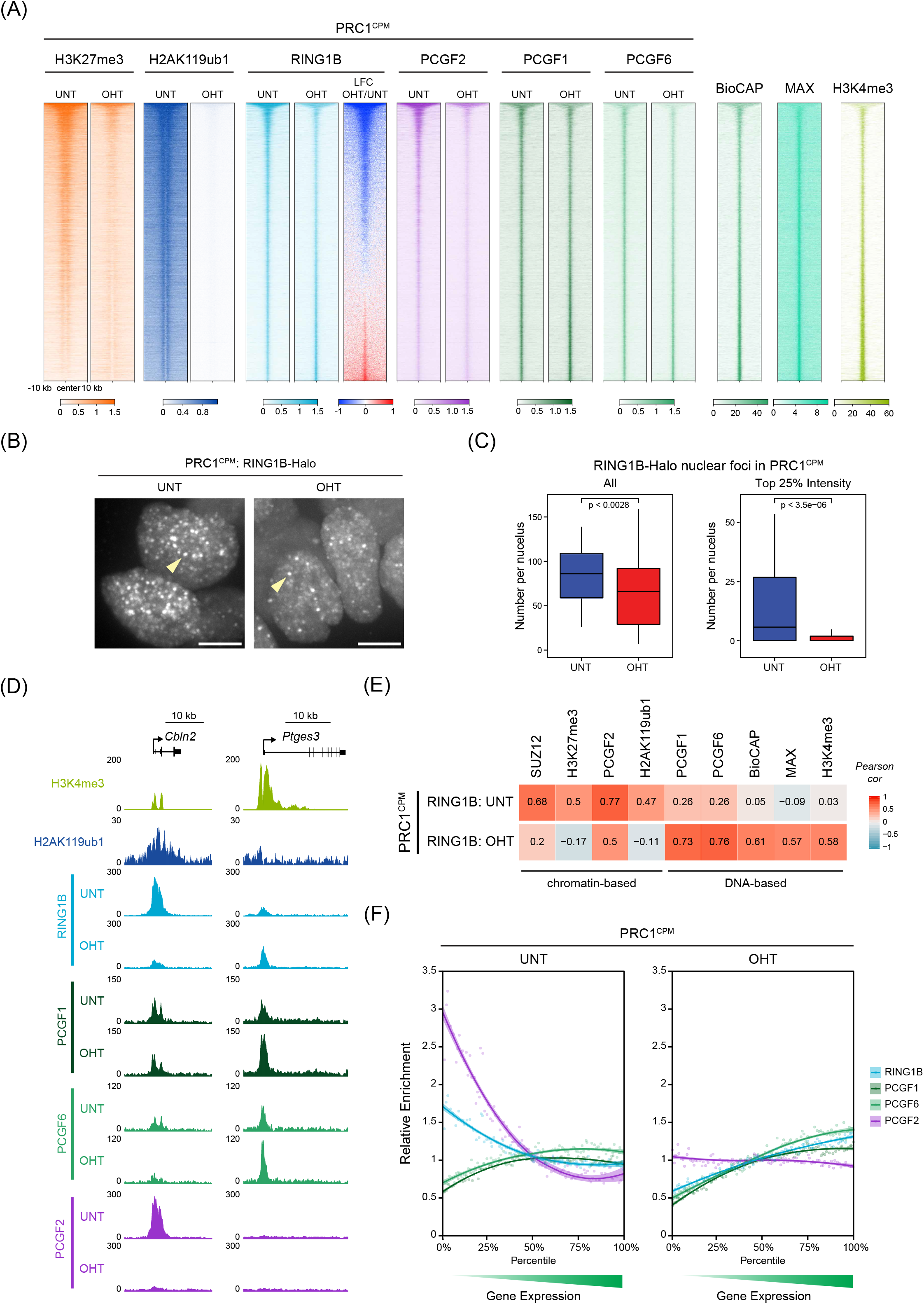
Removal of PRC1 catalysis reveals DNA-based sampling of target sites by vPRC1. **(A)** Heatmap analyses of cChIP-seq for H3K27me3, H2AK119ub1, RING1B, PCGF2, PCGF1 and PCGF6 at RING1B-bound sites in PRC1^CPM^ cells (untreated and OHT-treated). Also shown is BioCAP-seq (measure of non-methylated CpG-rich DNA) and ChIP-seq for MAX and H3K4me3 in wild-type ESCs. The genomic intervals were sorted based on the log2 fold change in RING1B occupancy following tamoxifen treatment in PRC1^CPM^ ESCs. **(B)** Maximum intensity projections of RING1B-Halo-JF_549_ signal in PRC1^CPM^ ESCs (untreated and OHT-treated). Examples of RING1B nuclear foci (Polycomb bodies) are indicated by arrowheads. Scale bar is 5 µm. **(C)** Box plots comparing the number of all (*left panel*) or top 25% highest intensity RING1B-Halo-JF_549_ foci per nucleus (*right panel*) in PRC1^CPM^ ESCs before (n_cells_ = 69) and after OHT treatment (n_cells_ = 83). P-values denote statistical significance calculated by a Student’s t-test. **(D)** Genomic snapshots of two genes, showing cChIP-seq for RING1B, PCGF1, PCGF6 and PCGF2 in PRC1^CPM^ ESCs (untreated and OHT-treated). H2AK119ub1 cChIP-seq in untreated cells and H3K4me3 ChIP-seq in wild-type ESCs is also shown. *Cbln2* is a lowly transcribed gene with high-level RING1B occupancy which relies on PRC1 catalysis and *Ptges3* is a more highly transcribed gene at which low-level RING1B binding is independent of catalysis. **(E)** Correlation of RING1B cChIP-seq signal in untreated and OHT-treated PRC1^CPM^ ESCs with cChIP-seq signal of PRC2 (SUZ12 and H3K27me3) and PRC1 (PCGF2, PCGF1, PCGF6, and H2AK119ub1) in untreated and OHT-treated PRC1^CPM^ ESCs. Correlation with BioCAP-seq and ChIP-seq for MAX and H3K4me3 in wild-type ESCs is also shown. This reveals a switch from chromatin-based to DNA-based target site selection by PRC1 following loss of PRC1 catalysis. **(F)** Relative enrichment of RING1B, PCGF1, PCGF6 and PCGF2 cChIP-seq signal in PRC1^CPM^ ESCs (untreated and OHT-treated) across promoter-proximal RING1B-bound sites divided into percentiles based on the expression level of the associated gene in untreated PRC1^CPM^ cells. For each factor, enrichment is shown relative to the fiftieth percentile. Lines represent smoothed conditional means based on loess local regression fitting. See also Figure S4.

To understand the mechanisms defining RING1B targeting in the absence of PRC1 catalytic activity, we turned our attention to the PCGF1- and PCGF6-containing vPRC1 complexes, both of which incorporate DNA binding modules. The PCGF1-PRC1 complex contains KDM2B, a ZF-CXXC domain protein that binds non-methylated CpGs (Farcas et al., 2012; He et al., 2013; Wu et al., 2013), while PCGF6-PRC1 incorporates MAX/MGA DNA binding factors (Gao et al., 2012; Hauri et al., 2016; Kloet et al., 2016; Ogawa et al., 2002). In untreated PRC1^CPM^ cells, cChIP-seq for PCGF1 and PCGF6 revealed a broad and generally uniform binding of both factors across all RING1B-occupied sites, which correlated poorly with RING1B levels (Figure 6A, D and E, Figure S4A and G). Importantly, in the tamoxifen-treated PRC1^CPM^ cells, we observed a strong correlation of RING1B occupancy with binding of PCGF1 and PCGF6, as well as with non-methylated CpG-rich DNA and MAX levels (Figure 6A and E and S4G and H). Furthermore, occupancy of PCGF1 and PCGF6 was only modestly affected by loss of PRC1 catalytic activity, indicating that binding of these vPRC1 complexes is largely independent and upstream of PRC1 catalysis and Polycomb chromatin domain formation (Figure 6A and D, Figure S4A and B). These observations strongly suggest that DNA binding activities inherent to PCGF1- and PCGF6-containing vPRC1 complexes constitute the primary targeting mechanisms that underpin target site selection by PRC1.

We and others have previously proposed that transcription may directly or indirectly counteract the formation of Polycomb chromatin domains (Jermann et al., 2014; Klose et al., 2013; Riising et al., 2014). In support of this idea, in untreated PRC1^CPM^ cells, the highest levels of RING1B and PCGF2 were associated with promoters of lowly transcribed genes that were enriched for H3K27me3 and H2AK119ub1, but not H3K4me3, a histone modification enriched at transcribed genes (Figure 6A and F, Figure S4I and J). In contrast, more actively transcribed RING1B targets exhibited low levels of H2AK119ub1, PRC2, H3K27me3 and cPRC1, despite being occupied by similar levels of the PCGF1 and PCGF6 vPRC1 complexes (Figure 6A and F, Figure S4I and J). Importantly, in tamoxifen-treated PRC1^CPM^ cells, strong enrichment of RING1B at lowly transcribed genes was lost, uncovering an underlying and largely uniform distribution of RING1B across target sites which mirrored the occupancy PCGF1 and PCGF6 (Figure 6A, D and F, Figure S4I and J). Together these observations are consistent with a model in which the PCGF1- and PCGF6 vPRC1 complexes broadly engage with, or sample, Polycomb target sites via their DNA binding domains, which, in the absence of counteracting features associated with active transcription, can lead to high-level H2AK119ub1 deposition and subsequent Polycomb chromatin domain formation.

### The catalytic activity of PRC1 is required for Polycomb-mediated gene repression

PRC1 is essential for Polycomb chromatin domain formation and this relies on catalysis (Figures 4-6), but whether PRC1 catalytic activity is also required for transcriptional repression still remains unresolved. To directly address this fundamental question, we carried out calibrated RNA-seq (cRNA-seq) in the isogenic PRC1^CPM^ and PRC1^CKO^ ESCs before and after tamoxifen treatment. Loss of PRC1 in PRC1^CKO^ cells resulted in a dramatic derepression of Polycomb target genes (Figure 7A and S5A), which is consistent with the central role of PRC1 in Polycomb-mediated gene repression in ESCs (Endoh et al., 2008; Fursova et al., 2019). Strikingly, removal of PRC1 catalysis resulted in reactivation of a similar number of Polycomb target genes (Figure 7A and Figure S5A), indicating that the catalytic activity of PRC1 is essential for Polycomb-mediated gene repression.

**Figure 7.**
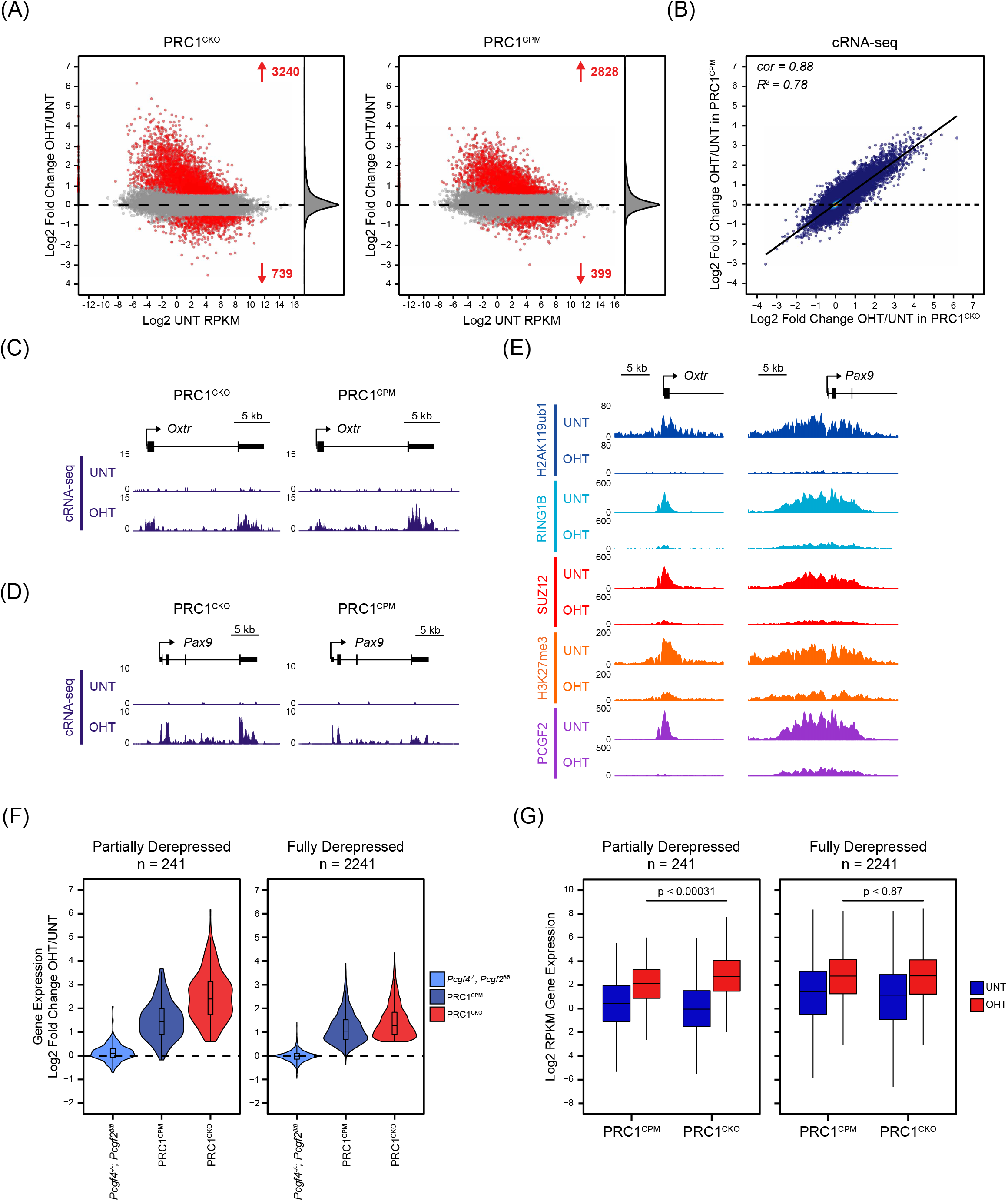
The catalytic activity of PRC1 is required for PRC1-mediated gene repression. **(A)** MA-plots of log2 fold changes in gene expression (cRNA-seq) in PRC1^CKO^ and PRC1^CPM^ ESCs following OHT treatment. Significant gene expression changes (p-adj < 0.05 and > 1.5-fold) are shown in red. Density of gene expression changes is shown on the right. **(B)** A scatter plot comparing the log2 fold changes in gene expression (cRNA-seq) in PRC1^CKO^ and PRC1^CPM^ ESCs following OHT treatment for all genes. *R^2^* represents coefficient of determination for linear regression, and *cor* denotes Pearson correlation coefficient. **(C)** A genomic snapshot of a PRC1-repressed gene which is fully derepressed in OHT-treated PRC1^CPM^ as compared to PRC1^CKO^ ESCs, showing gene expression (cRNA-seq) in PRC1^CKO^ and PRC1^CPM^ ESCs (untreated and OHT-treated). **(D)** A genomic snapshot of a PRC1-repressed gene which is partially derepressed in OHT-treated PRC1^CPM^ as compared to PRC1^CKO^ ESCs, showing gene expression (cRNA-seq) in PRC1^CKO^ and PRC1^CPM^ ESCs (untreated and OHT-treated). **(E)** Genomic snapshots for the genes described in (C-D), showing cChIP-seq for PRC1 (RING1B, H2AK119ub1 and PCGF2) and PRC2 (SUZ12 and H3K27me3) in PRC1^CPM^ ESCs (untreated and OHT-treated). **(F)** A violin plot comparing log2 fold changes in expression (cRNA-seq) of PRC1-repressed genes which are partially (n = 241) or fully derepressed (n = 2241) in PRC1^CPM^ as compared to PRC1^CKO^ ESCs following OHT treatment in *Pcgf4^-/-^;Pcgf2^fl/fl^*, PRC1^CPM^ and PRC1^CKO^ ESC lines. **(G)** A box plot of gene expression levels (cRNA-seq) for genes that are partially (n = 241) or fully derepressed (n = 2241) in PRC1^CPM^ as compared to PRC1^CKO^ ESCs following OHT treatment in PRC1^CPM^ and PRC1^CKO^ ESCs (untreated and OHT-treated). P-values denote statistical significance calculated by a Wilcoxon signed-rank test. See also Figure S5.

A more detailed comparison of the gene expression alterations in the tamoxifen-treated PRC1^CKO^ and PRC1^CPM^ ESCs revealed that the magnitude of gene expression changes strongly correlated between these cell lines, both when looking at all genes and specifically at PRC1-repressed genes (Figure 7B and S5B and C). This demonstrates that loss of PRC1 catalytic activity largely phenocopies the gene expression defects that manifest when PRC1 is completely removed. However, we found a small number of PRC1-repressed genes (241 out of 2482) which appeared to be derepressed to a lesser extent in the PRC1^CPM^ compared to the PRC1^CKO^ ESCs following tamoxifen treatment (Figure 7D, F and G). These tended to have large Polycomb chromatin domains that were highly enriched with cPRC1 and H3K27me3 in untreated cells and retained low level of cPRC1 binding in the tamoxifen-treated PRC1^CPM^ cells (Figure 7E and Figure S5D and E). Nevertheless, repression of this small subset of genes still relied heavily on PRC1 catalysis, as removal of cPRC1 alone caused only a modest increase in their expression (Figure 7F). Importantly, the remaining PRC1-repressed genes were reactivated to the same level in PRC1^CPM^ and PRC1^CKO^ ESCs following tamoxifen treatment (Figure 7C and G), indicating that PRC1 catalysis is a central determinant of gene repression. Therefore, through systematically dissecting the requirement for PRC1 catalytic activity in Polycomb system function, we provide direct evidence that this activity is essential for both Polycomb chromatin domain formation and gene repression.

## Discussion

The Polycomb system represents a paradigm for chromatin-based gene regulation. However, the mechanisms by which Polycomb target sites are identified and repressive Polycomb chromatin domains are formed have remained elusive. Moreover, the extent to which this relies on PRC1 catalysis is still controversial. Here, using *in vitro* enzymatic assays and genome engineering in ESCs, we discover that a mutation (RING1B^I53A^) previously used to study the role of PRC1 catalysis in Polycomb system function, is not catalytically inactive (Figures 1 and 2). We develop an elegant conditional strategy to completely ablate PRC1 catalytic activity while leaving PRC1 complexes intact (Figure 3). Using this new tool, we discover that catalysis by PRC1 is essential for PRC2 occupancy and H3K27me3 deposition at target sites, together with downstream recruitment of cPRC1 complexes which support long-range interactions between Polycomb chromatin domains (Figure 4-5). We show that DNA-binding vPRC1 complexes broadly occupy target sites independently of PRC1 catalysis and that this likely represents a primary mechanism of Polycomb target site selection prior to Polycomb chromatin domain formation (Figure 6). Finally, and most importantly, we demonstrate that the catalytic activity of PRC1 is essential for Polycomb-mediated gene repression (Figure 7). Together, these discoveries demonstrate a central requirement for PRC1 catalysis in gene repression and provide compelling new evidence for a PRC1-initiated pathway that drives Polycomb chromatin domain formation and gene repression.

Understanding whether catalysis by chromatin modifying enzymes is central to their function is an ongoing challenge in the field of chromatin biology. Definitive interpretations can only be arrived at when *in vitro* biochemical characterisations demonstrate that inactivating mutations cause complete loss of catalysis without protein complex disruption. Furthermore, this must be combined with quantitative measurements of the modified substrate *in vivo*, to ensure that the product of catalysis is completely lost in the cellular context. Here we satisfy these two central requirements for studying PRC1 catalytic activity, whose contribution to Polycomb system function has remained controversial. Importantly, we discover that catalysis by PRC1 is essential for normal Polycomb chromatin domain formation and gene repression. While we have limited our investigation to ESCs, where the Polycomb system has been extensively characterised, a recent study has also examined the contribution of PRC1 catalytic activity to gene regulation using a neuronal cell fate restriction model (Tsuboi et al., 2018). In agreement with our observations in ESCs, it was reported that RING1B^I53A^ mutation is hypomorphic, while RING1B^I53A/D56K^ is catalytically inactive. Furthermore, from expression analysis of a small number of neurogenic genes, this study concluded that PRC1 catalytic activity is required for gene repression during the early stages of neurogenic commitment. Interestingly, however, at later stages of neuronal lineage specification, gene repression did not appear to rely solely on PRC1 catalysis. This suggests that catalysis-independent mechanisms may contribute to the maintenance of gene repression in more slowly- or non-dividing cells that are characteristic of later developmental stages, as has been previously proposed (Cohen et al., 2018). Moving forward, it will be important to use the catalytic mutations we have extensively characterised to examine at the genome scale how widely the enzymatic activity of PRC1 contributes to Polycomb system function during mammalian development. Furthermore, these mutations should also be used in other model organisms, such as *Drosophila*, where the contribution of PRC1 catalysis to gene regulation and development has been proposed to be more limited (Pengelly et al., 2015).

Defining the mechanisms of Polycomb target site selection and repressive Polycomb chromatin domain formation is central to understanding the logic by which the Polycomb system functions. In mammalian cells, high-level Polycomb occupancy occurs at CpG island (CGI) elements (Mikkelsen et al., 2007), while ectopic GC-rich DNA is sufficient to establish a new Polycomb chromatin domain *de novo* (Jermann et al., 2014; Lynch et al., 2012; Mendenhall et al., 2010). A mechanistic link between CGIs and Polycomb recruitment came with the discovery that KDM2B, a component of the PCGF1-containing vPRC1 complex, has DNA binding activity which is specific for non-methylated CpG dinucleotides, suggesting that recognition of CGI DNA may underpin Polycomb target site selection (Farcas et al., 2012; He et al., 2013; Wu et al., 2013). Similarly, the PCGF6-containing vPRC1 complex has DNA binding activities that contribute to Polycomb occupancy in specialised contexts (Endoh et al., 2017; Zdzieblo et al., 2014) or more generally (Fursova et al., 2019; Scelfo et al., 2019; Yang et al., 2016). However, while PCGF1 and PCGF6 broadly occupy target sites, somewhat paradoxically only a subset of these achieve high levels of PRC1, H2AK119ub1, PRC2 and H3K27me3 (Figure 6). Here, using our PRC1^CPM^ ESCs, we provide new evidence for a model in which the PCGF1- and PCGF6-containing vPRC1 complexes utilise their DNA binding activities to sample all potential Polycomb target genes, but high-level deposition of H2AK119ub1 and subsequent Polycomb chromatin domain formation occurs only at target genes with low transcriptional activity. This model is supported by previous work showing that the Polycomb system responds to, as opposed to instructs, the transcriptional state of a gene (Berrozpe et al., 2017; Hosogane et al., 2013; Riising et al., 2014). We propose that at lowly transcribed genes a PRC1-initiated pathway that relies on catalysis is required to protect against low-level stochastic gene activation events that would otherwise be deleterious to maintenance of cell type-specific gene expression programmes.

Our observations reveal that catalysis by PRC1 is essential for Polycomb-mediated gene repression, but how is this achieved mechanistically? We recently demonstrated that removal of cPRC1 complexes in ESCs has little effect on Polycomb-mediated gene repression (Fursova et al., 2019), while earlier work showed that PRC2 removal and loss of H3K27me3 resulted in few gene expression defects (Riising et al., 2014), suggesting that these activities may play a secondary role and instead contribute to the fidelity of gene repression. Based on these findings, it seems unlikely that the role of PRC1 catalysis and H2AK119ub1 is simply to recruit PRC2, H3K27me3 and cPRC1 complexes to elicit gene repression. One possibility is that H2AK119ub1 could directly disrupt RNA Polymerase II activity (Stock et al., 2007; Zhou et al., 2008), as recently suggested by *in vitro* transcription assays, in which incorporation of H2AK119ub1 into recombinant chromatin templates blocked transcription (Aihara et al., 2016; Nakagawa et al., 2008). Alternatively, PRC1 catalytic activity and H2AK119ub1 could have an indirect effect on transcription, for example by creating a binding site for reader proteins which would then elicit repression (Ali et al., 2018; Cooper et al., 2016; Qin et al., 2015; Richly et al., 2010; Zhang et al., 2017). In addition, incorporation of a bulky ubiquitin moiety may interfere with deposition of other histone modifications that facilitate gene expression (Nakagawa et al., 2008; Yuan et al., 2013). Finally, the possibility that PRC1 drives gene repression via ubiquitylation of histone H2A variants (Surface et al., 2016) or other non-histone substrates cannot be ruled out (Ben-Saadon et al., 2006). Clearly, future work focussed on defining the mechanisms by which the catalytic activity of PRC1 counteracts RNA Polymerase II activity and gene expression will be important. Nevertheless, our new discoveries place PRC1 catalysis at the forefront of Polycomb-mediated gene repression, paving the way for a mechanistic understanding of Polycomb function.

## Acknowledgements

Work in the Klose lab is supported by the Wellcome Trust, the European Research Council, and the Lister Institute of Preventive Medicine. We would like to thank Amanda Williams at the Department of Zoology, Oxford, for sequencing support on the NextSeq 500. We are grateful to Chloe Roustan and Neil Brockdorff for sharing bacterial expression constructs for PRC1. We thank Paula Dobrinić and Emma Smith for critical reading of the manuscript.

## Author Contributions

Conceptualization, N.P.B, N.A.F., and R.J.K.; Methodology, N.P.B., N.A.F., J.R.K., M.K.H., and A.F.; Investigation, N.P.B, N.A.F., J.R.K., M.K.H., and A.F.; Formal analysis, N.P.B., N.A.F., J.R.K., M.K.H., and A.F.; Resources, N.P.B, N.A.F., and M.K.H.; Writing – Original Draft, N.P.B., N.A.F., and R.J.K.; Writing –Review and Editing, N.P.B., N.A.F., J.R.K., M.K.H., A.F., and R.J.K.; Funding acquisition, R.J.K.; Supervision, R.J.K.

---

**Figure S1. Related to Figure 3.**
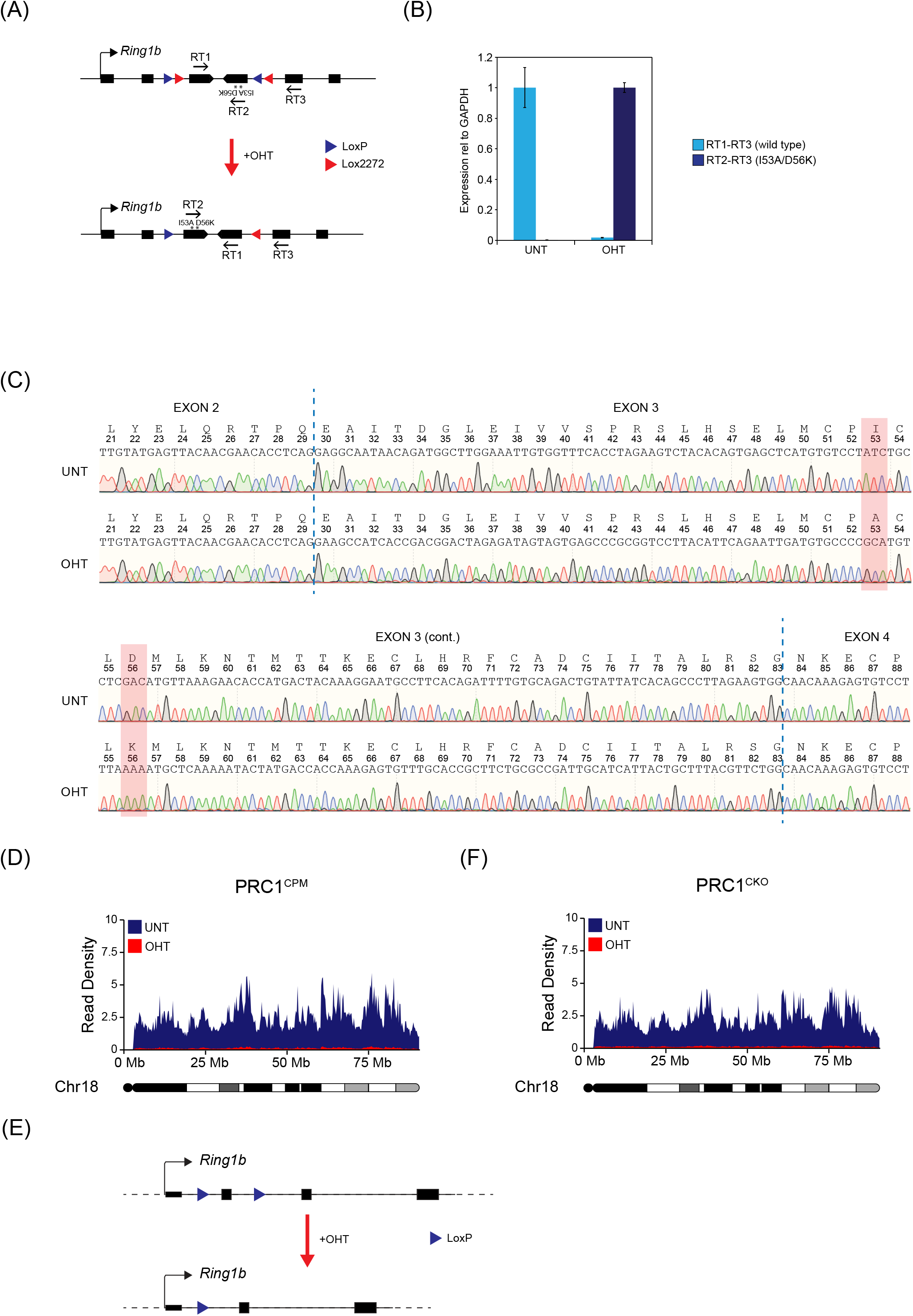
**(A)** A schematic of the endogenous *Ring1b* allele in PRC1^CPM^ cell line before and after addition of OHT, showing positions of the primers used for RT-qPCR quantification of conversion from *Ring1b^WT^*to *Ring1b^I53A/D56K^*. In untreated cells, incorporation of wild-type exon 3 into *Ring1b* mRNA gives RT-qPCR signal from the primer pair RT1-RT3, but not RT1-RT2. Following OHT treatment and flipping of the exon 3 cassette, incorporation of I53A/D56K version of exon 3 into *Ring1b* mRNA gives RT-qPCR signal from the primer pair RT2-RT3, but not RT1-RT3. **(B)** RT-qPCR validation of PRC1^CPM^ line using primers described in (A). Error bars represent SEM (n=4). **(C)** DNA sequencing traces of *Ring1b* mRNA in untreated and OHT-treated PRC1^CPM^ cells, showing complete conversion from RING1B^WT^ to RING1B^I53A/D56K^. Vertical dashed lines indicate boundaries between exons. Amino acid sequences are shown above corresponding DNA sequence, with I53A and D56K positions highlighted in red. Wild-type and I53A/D56K mutant versions of exon 3 were engineered to be different at wobble position of each triplet codon, to allow each exon to be easily distinguished and minimize formation of secondary RNA structures that could impact on splicing. **(D)** A chromosome density plot showing H2AK119ub1 cChIP-seq across chromosome 18 in untreated (blue) and OHT-treated (red) PRC1^CPM^ cells. **(E)** A schematic of the endogenous *Ring1b* allele in PRC1^CKO^ cell line before and after addition of OHT, showing parallel LoxP sites flanking exon 2 (the first coding exon). OHT treatment causes CRE-mediated deletion of exon 2 which puts the rest of the *Ring1b* coding sequence out of frame, resulting in no functional protein being produced. **(F)** A chromosome density plot showing H2AK119ub1 cChIP-seq data across chromosome 18 in untreated (blue) and OHT-treated (red) PRC1^CKO^ cells.

**Figure S2. Related to Figure 4.**
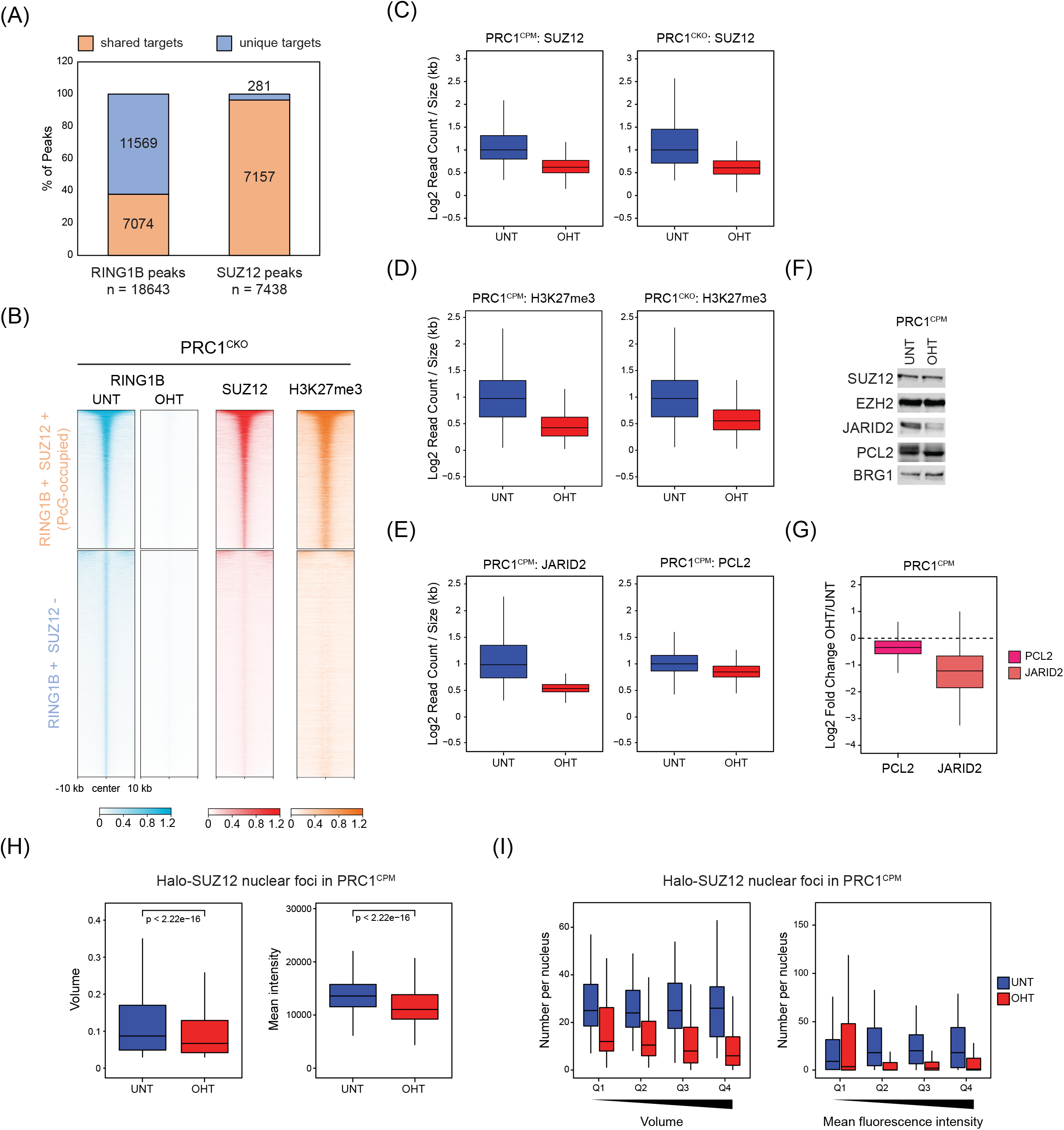
**(A)** Bar plots showing the overlap between RING1B- and SUZ12-bound genomic regions. This illustrates that PRC1 and PRC2 share a large number of target sites, with > 95% of SUZ12 peaks overlapping with RING1B peaks, while RING1B binding appears to be more broad. **(B)** Heatmap analysis of cChIP-seq in PRC1^CKO^ ESCs for RING1B (untreated and OHT-treated), SUZ12 and H3K27me3 (untreated) at RING1B-bound sites divided into two groups based on the overlap with SUZ12 peaks (RING1B/SUZ12-bound or PcG-occupied (n = 7074) and only RING1B-bound (n = 11,569)). For each group, the genomic regions were sorted based on RING1B occupancy in untreated cells. **(C)** Box plots of the normalised cChIP-seq signal for SUZ12 at PcG-occupied sites for untreated and OHT-treated PRC1^CPM^ and PRC1^CKO^ cells. **(D)** Box plots of the normalised cChIP-seq signal for H3K27me3 at PcG-occupied sites for untreated and OHT-treated PRC1^CPM^ and PRC1^CKO^ cells. **(E)** Box plots of the normalised cChIP-seq signal for JARID2 and PCL2 at PcG-occupied sites for untreated and OHT-treated PRC1^CPM^ cells. **(F)** Western blot analysis of PRC2 subunits (SUZ12, EZH2, JARID2 and PCL2) in untreated and OHT-treated PRC1^CPM^ ESCs. BRG1 was used as a loading control. **(G)** Box plot of log2 fold changes in PCL2 and JARID2 binding at PcG-occupied sites following OHT treatment in PRC1^CPM^ cells. **(H)** Box plots comparing volumes (*left*) and mean intensities (*right*) of JF_549_-Halo-SUZ12 nuclear foci in PRC1^CPM^ ESCs before (n_cells_ = 55) and after OHT treatment (n_cells_ = 52). P-values denote statistical significance calculated by a Wilcoxon signed-rank test. **(I)** Box plots comparing numbers of JF_549_-Halo-SUZ12 nuclear foci detected in PRC1^CPM^ ESCs before (n_ells_ = 55) and after OHT treatment (n_cells_ = 52), with foci divided into quartiles based on foci volume (*left*) or mean intensities (*right*) in untreated cells (from lowest Q1 to highest Q4). This shows that in the absence of PRC1 catalysis, SUZ12 nuclear foci of different volumes and intensities are lost, with largest and brightest foci being affected the most.

**Figure S3. Related to Figure 5.**
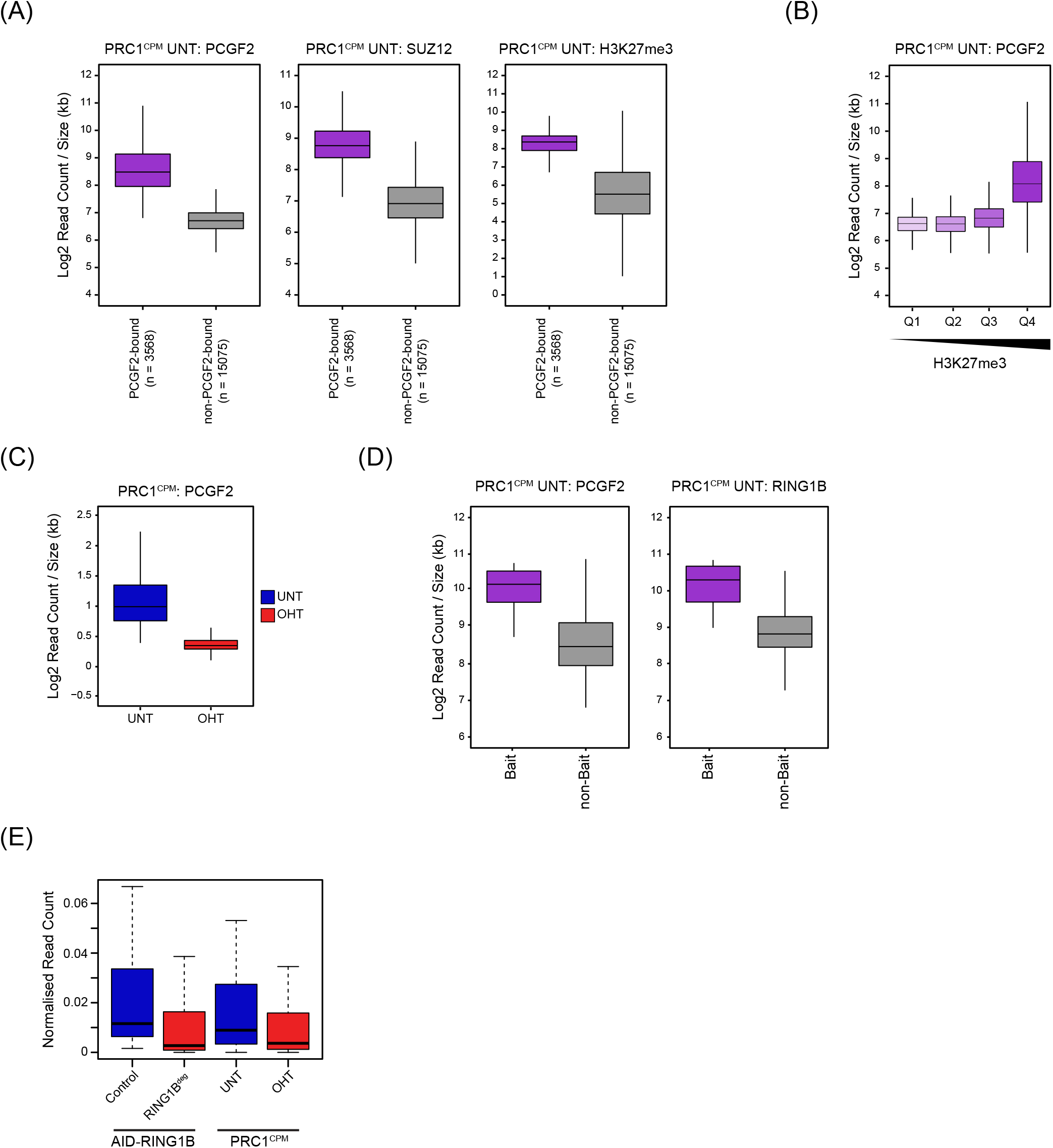
**(A)** Box plot of the cChIP-seq signal for PCGF2, SUZ12 and H3K27me3 in untreated PRC1^CPM^ cells at RING1B-bound sites divided into sites that are bound (PCGF2 target sites) or not bound by PCGF2. **(B)** Box plots comparing the cChIP-seq signal for PCGF2 in untreated PRC1^CPM^ cells at RING1B-bound sites divided into quartiles (from lowest Q1 to highest Q4) based on H3K27me3 levels. **(C)** Box plots of the normalised cChIP-seq signal for PCGF2 at PCGF2 target sites in untreated and OHT-treated PRC1^CPM^ ESCs. **(D)** Box plots comparing PCGF2 and RING1B cChIP-seq signal in untreated PRC1^CPM^ cells at PCGF2 target sites used as baits in CaptureC analysis and the remaining PCGF2 target sites. **(E)** Box plot analysis of the normalised mean read count from CaptureC for the interactions between the bait Polycomb target regions and other PCGF2 target sites (n = 130) in auxin-treated *Ring1a^-/-^*;AID-RING1B (RING1B^deg^) ESCs (relative to the Control cell line with intact PRC1) and PRC1^CPM^ cells (untreated and OHT-treated).

**Figure S4. Related to Figure 6.**
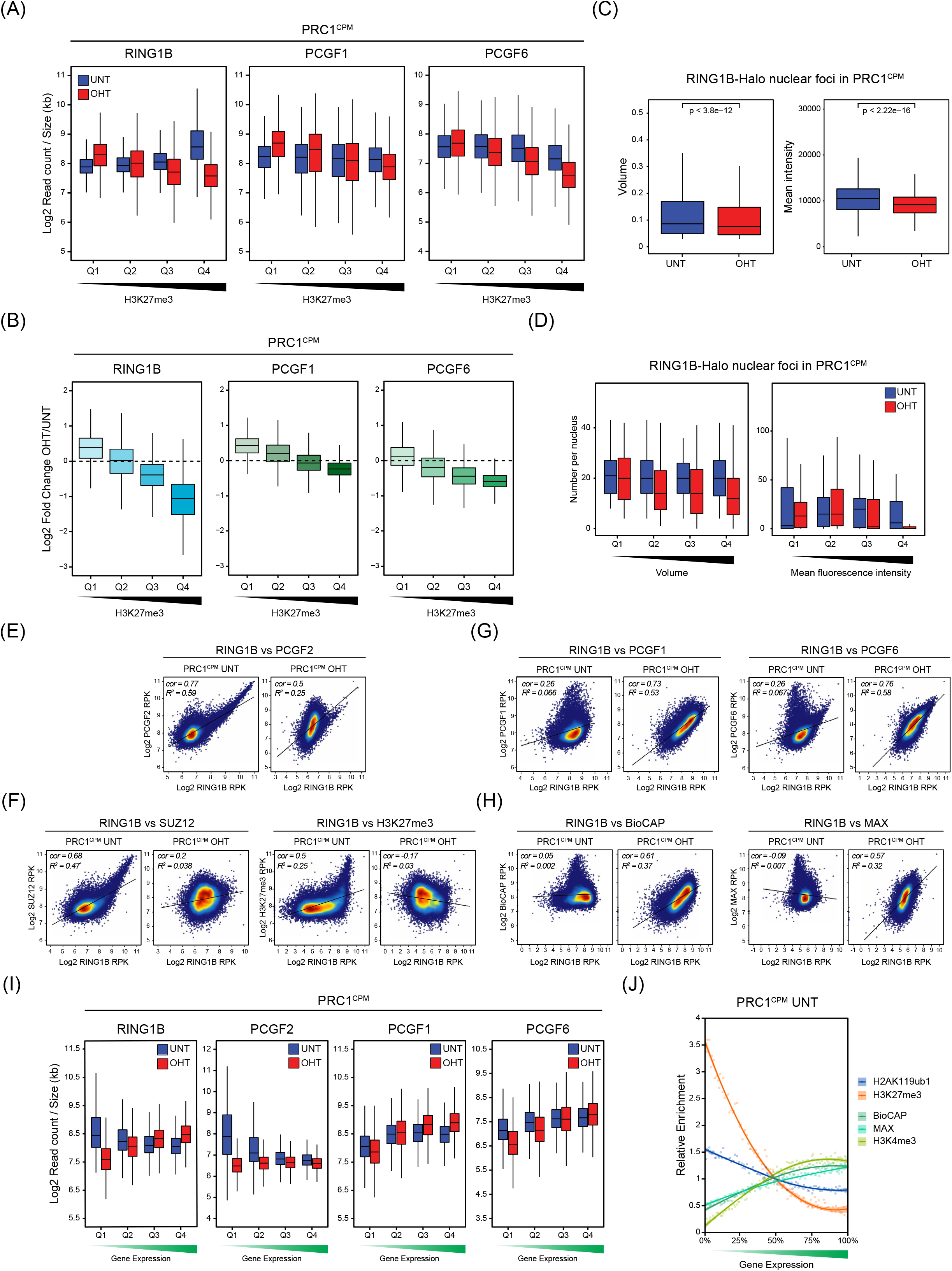
**(A)** Box plots of cChIP-seq signal for RING1B, PCGF1 and PCGF6 at RING1B-bound sites divided into quartiles based on H3K27me3 levels in untreated cells (from lowest Q1 to highest Q4) in PRC1^CPM^ ESCs before (UNT, blue) and after OHT treatment (OHT, red). This shows that PCGF1 and PCGF6 occupancy is largely uniform across all RING1B-bounds sites with different H3K27me3 levels. **(B)** Box plots comparing changes in cChIP-seq signal for RING1B, PCGF1 and PCGF6 following OHT treatment in PRC1^CPM^ cells at RING1B-bound sites divided into quartiles based on H3K27me3 levels in untreated cells. Together with (A), this shows that following removal of PRC1 catalytic activity, RING1B occupancy is reduced at sites with high levels of H3K27me3 (Q3 and Q4) but is retained/increased at sites with low levels of H3K27me3 (Q1 and Q2). In addition, it demonstrates that PCGF1 and PCGF6 occupancy is only modestly affected following loss of PRC1 catalytic activity, with binding of PCGF1 slightly increased at sites with low level of H3K27me3 and binding of PCGF6 moderately reduced at sites with high enrichment of H3K27me3. **(C)** Box plots comparing volumes and mean intensities of RING1B-Halo-JF_549_ nuclear foci in PRC1^CPM^ ESCs before (n_cells_ = 69) and after OHT treatment (n_cells_ = 83). P-values denote statistical significance calculated by a Wilcoxon signed-rank test. **(D)** Box plots comparing numbers of RING1B-Halo-JF_549_ nuclear foci detected in PRC1^CPM^ ESCs before (n_cells_ = 69) and after OHT treatment (n_cells_ = 83), with foci divided into quartiles based on foci volume (*left*) or mean intensities (*right*) in untreated cells (from lowest Q1 to highest Q4). This shows that loss of PRC1 catalysis had the strongest effect on RING1B nuclear foci with the largest volume and mean intensity, while the foci with low to moderate intensities/volumes remained largely unchanged or even increased. This is in agreement with changes in RING1B occupancy by cChIP-seq at target sites with different levels of RING1B binding in untreated cells. **(E)** Scatter plots showing the relationship between the cChIP-seq signals of RING1B and PCGF2 at RING1B-bound sites in PRC1^CPM^ ESCs before (UNT) and after OHT treatment (OHT). *R^2^* represents coefficient of determination for linear regression and *cor* denotes Pearson correlation coefficient. **(F)** As in (E) for RING1B and SUZ12 cChIP-seq (*left*) and RING1B and H3K27me3 cChIP-seq (*right*). **(G)** As in (E) for RING1B and PCGF1 cChIP-seq (*left*) and RING1B and PCGF6 cChIP-seq (*right*). **(H)** Scatter plots showing the relationship between the cChIP-seq signal for RING1B at RING1B-bound sites in PRC1^CPM^ ESCs before (UNT) and after OHT treatment (OHT) with BioCAP-seq (*left*) or MAX ChIP-seq signal (*right*) in wild-type ESCs. *R^2^* represents coefficient of determination for linear regression and *cor* denotes Pearson correlation coefficient. **(I)** Box plots of RING1B, PCGF2, PCGF1 and PCGF6 cChIP-seq signal in untreated (UNT, blue) and OHT-treated (OHT, red) PRC1^CPM^ ESCs at promoter-proximal RING1B-bound sites divided into quartiles based on the expression level of the associated gene in untreated cells (from lowest Q1 to highest Q4). **(J)** Relative enrichment of H2AK119ub1 and H3K27me3 cChIP-seq signal in untreated PRC1^CPM^ ESCs and H3K4me3 and MAX ChIP-seq, as well as BioCAP-seq signal in wild-type ESCs across promoter-proximal RING1B-bound sites divided into percentiles based on the expression level of the associated gene in untreated PRC1^CPM^ cells. For each factor, enrichment is shown relative to the fiftieth percentile. Lines represent smoothed conditional means based on loess local regression fitting.

**Figure S5. Related to Figure 7.**
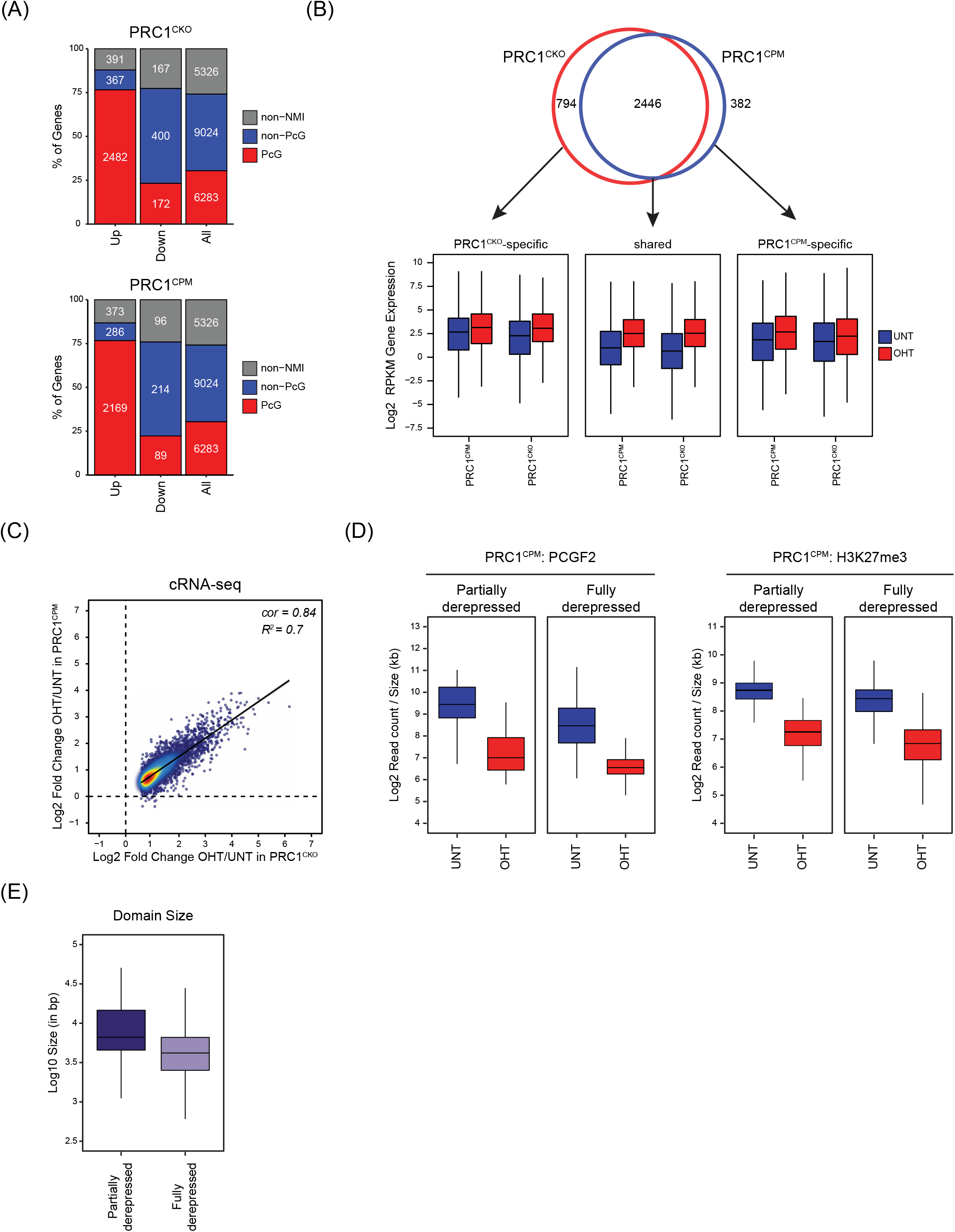
**(A)** A bar plot illustrating the distribution of gene expression changes (p-adj < 0.05 and > 1.5-fold) in cRNA-seq between different gene classes (Non-NMI, Non-PcG, PcG) for PRC1^CKO^ and PRC1^CPM^ ESCs following OHT treatment. Non-NMI genes (n = 5326) lack a non-methylated CpG island (NMI) at their promoter. Non-Polycomb target genes (n = 9024) have an NMI promoter that is not co-occupied by both Polycomb repressive complexes. Polycomb target genes (n = 6283) have an NMI promoter that is bound by both PRC1 and PRC2. **(B)** *Top panel:* A Venn diagram showing the overlap between the genes derepressed (p-adj < 0.05 and > 1.5-fold) in PRC1^CKO^ (red) and PRC1^CPM^ (blue) ESCs following OHT treatment. *Bottom panel:* Box plots comparing the gene expression levels from cRNA-seq in PRC1^CPM^ and PRC1^CKO^ ESCs before (UNT) and after OHT treatment (OHT) for three groups of derepressed genes defined in the Venn diagram above: PRC1^CKO^-specific, shared between PRC1^CKO^ and PRC1^CPM^ ESCs, and PRC1^CPM^-specific. This demonstrates that for the genes categorised as specifically derepressed in either PRC1^CKO^ or PRC1^CPM^ OHT-treated ESCs, the magnitude of gene expression changes is more modest than for the shared targets, making them more sensitive to the choice of a significance threshold. Importantly, expression levels of these genes following OHT treatment are highly similar between the two cell lines. **(C)** A scatter plot comparing the log2 fold changes in gene expression in cRNA-seq following OHT treatment in PRC1^CPM^ and PRC1^CKO^ ESCs for PRC1-repressed genes. *R^2^* represents coefficient of determination for linear regression and *cor* denotes Pearson correlation coefficient. **(D)** Box plots of PCGF2 cChIP-seq signal at the RING1B-bound sites associated with the promoters of PRC1-repressed genes which are either partially (n = 241) or fully derepressed (n = 2241) following loss of PRC1 catalysis as compared to a complete loss of PRC1 in PRC1^CPM^ cells before (UNT) and after OHT treatment (OHT). **(E)** As in (E) for H3K27me3 cChIP-seq. **(F)** Box plots comparing sizes of RING1B-bound sites overlapping with the promoters of PRC1-repressed genes which are either partially (n = 241) or fully derepressed (n = 2241) following OHT treatment in PRC1^CPM^ as compared to PRC1^CKO^ ESCs.

## Methods

### Recombinant protein expression and purification

Mouse RING1B, tagged with StrepII and 6xHis tags, was coexpressed with PCGF1 and RYBP from a pST44 polycistronic plasmid in *E. coli* BL21 (DE3) pLysS. Cultures were supplemented during expression with 250 µM ZnCl_2_. Cells were lysed by sonication in lysis buffer containing 20 mM Tris (pH 8.0), 500 mM NaCl, 0.1% NP-40 and cOmplete Protease Inhibitor Cocktail (Roche) and trimeric complexes were affinity purified via 6xHis-tagged RING1B on Ni^2+^-charged IMAC Sepharose 6 Fast Flow resin (GE Healthcare). 10 mM imidazole was added to lysates during binding. Wash buffer contained 50 mM NaH_2_PO_4_ (pH 8.0), 300 mM NaCl and 20 mM imidazole and protein was eluted in wash buffer containing increasing imidazole (100-250 mM). Purified PRC1 complexes were dialysed into BC100 (50 mM HEPES (pH 7.9), 100 mM KCl, 10% Glycerol, 1 mM DTT).

### Reconstitution of Nucleosomes

Nucleosomes were reconstituted as described previously (Dyer et al., 2004). Recombinant *Xenopus* histones were expressed in *E. coli* BL21(DE3) pLysS and purified from inclusion bodies via Sephacryl S-200 gel filtration (GE Healthcare). Stoichiometric amounts of each core histone were incubated together under high salt conditions (2 M NaCl) and the resulting histone octamer purified using a Superdex 200 gel filtration column (GE Healthcare). Purified 147bp DNA carrying the 601 nucleosome-positioning sequence was a kind gift from the Brockdorff lab. Purified DNA, in slight excess to octamers, was mixed together in 2 M NaCl and diluted stepwise with 10 mM Tris (pH 7.5) to reach a final concentration of 100 mM NaCl. The reconstituted nucleosomes were analysed on a 0.8% Tris-borate agarose gel and concentrated using a 30,000 MWCO spin concentrator (GE Healthcare).

### E3 ubiquitin ligase assays

H2A ubiquitylation assays were carried out as previously (Rose et al., 2016). Briefly, UBE1 (Boston Biochem), UbcH5c (Enzo), methylated ubiquitin (Boston Biochem) and ATP (Life technologies) were pre-incubated for 20 min at 37°C prior to addition of reconstituted PRC1 and nucleosomes. Reactions were allowed to proceed for 1 hr at 37°C then quenched with 30 mM EDTA and subject to SDS-PAGE for western blot analysis. Western blots were probed with antibodies which recognise Histone H2A in both ubiquitylated and unmodified form (Millipore 07-146) and Histone H3 (CST, 96C10), followed by incubation with LiCOR IRDye secondary antibodies (800CW goat anti-rabbit and 680RD goat anti-mouse). Western blots were imaged using the LiCOR Odyssey Fc imaging system and band intensities were quantified using ImageStudio. H2A band intensities were normalised to H3 and the fraction of ubiquitylated H2A relative to total H2A was quantified. Data were visualised and dose-response curves fitted using GraphPad Prism 7.

### Genome engineering by CRISPR/Homology-Directed Repair (HDR)

The pSptCas9(BB)-2A-Puro(PX459)-V2.0 vector was obtained from Addgene (#62988) and sgRNAs were designed using the CRISPOR online tool (http://crispor.tefor.net/crispor.py). Targeting constructs with appropriate homology arms were generated by Gibson assembly using the Gibson Assembly Master Mix kit (New England Biolabs), or in the case of single LoxP sites with 150 bp homology arms, purchased from GeneArt Gene Synthesis (Invitrogen). In all instances, targeting constructs were designed such that Cas9 recognition sites were disrupted by the presence of the LoxP site. ESCs (one well of a 6-well plate) were transfected with 0.5 μg of each Cas9 guide, and 2 μg of targeting construct (where appropriate) using Lipofectamine 3000 (ThermoFisher) according to manufacturer’s guidelines. The day after transfection, cells were passaged at a range of densities and subjected to puromycin selection (1 μg/ml) for 48 hours to eliminate any non-transfected cells. Approximately one week later, individual clones were isolated, expanded, and PCR-screened for the desired genomic modification.

### Cell line generation

Constitutive *Ring1b^I53A^* and *Ring1b^I53A/D56K^*ESCs were generated from E14 ESCs using a Cas9 guide specific for the mutation site in the endogenous *Ring1b* gene, and targeting constructs with approximately 1 kb homology arms. The targeting constructs were designed so that as well as introducing the desired mutation into *Ring1b*, an *Msp*I restriction site was also created to enable screening of clones via a PCR and digest approach. Putative homozygote clones were carried forward for RT-PCR and sequencing to verify that the *Ring1b* transcript carried the desired mutation, as well as western blot analysis. An analogous strategy was used to insert the I50A/D53K mutation into *Ring1a* (see below).

To generate the PRC1^CPM^ line, a targeting construct was generated comprising exon 3 of *Ring1b* in forward orientation (flanked by 100 bp of *Ring1b* intron 2/intron 3) followed by a mutant copy of exon 3 (encoding I53A and D56K mutations) in reverse orientation (flanked by splice donor and acceptor sites from mouse *IgE* gene). Both the wild-type and mutant versions of exon 3 were codon optimized at wobble positions to minimize sequence similarity, thereby avoiding hairpin formation and allowing the two to be easily distinguished. This exon 3 pair was flanked by doubly inverted LoxP/Lox2272 sites and approximately 1 kb homology arms (see Supplementary Table 1 for sequence). To help ensure that the RING1B^CPM^ cassette was inserted correctly, the RING1B^CPM^ targeting construct was transfected into E14 ESCs in combination with three different Cas9 guides specific for the *Ring1b* locus (see Supplementary table 1). Correctly targeted homozygous clones were identified by PCR screening, followed by RT-PCR and sequencing to check for splicing defects. CreERT2 was then inserted into the *Rosa26* locus using a *Rosa26*-specific Cas9 guide, and using a similar approach, the I50A/D53K mutation was constitutively knocked into both copies of endogenous *Ring1a.* The final PRC1^CPM^ cell line was validated by PCR, RT-PCR and western blot, with and without tamoxifen treatment.

The isogenic PRC1^CKO^ line used in this study was described previously (Fursova et al., 2019). Briefly, exons 1-3 of *Ring1a* were firstly deleted using Cas9 guides flanking the 1.5 kb deletion region, and Cre-ERT2 was inserted into the *Rosa26* locus using a *Rosa26*-specific guide (see Supplementary Table 1). *Ring1a^-/-^;CreERT2* ESCs were then subjected to two sequential rounds of genome editing to insert parallel LoxP sites flanking exon 2 (the first coding exon) of *Ring1b*.

The control TIR1-only and AID-RING1B lines were generated from E14 ESCs as described previously (Rhodes et al., 2019). Briefly, Cas9 engineering was used to insert the coding sequence for *Oryza sativa* TIR1 into the *Rosa26* locus, thereby generating the TIR1-only control line. To generate the AID-RING1B line, the TIR1-only line was subjected to further rounds of Cas9-mediated engineering to introduce the auxin inducible degron (AID) tag at the N-terminus of both copies of *Ring1b*, and constitutively delete *Ring1a* using Cas9 guides flanking exons 1-3. Western blot analysis was used to confirm loss of RING1B protein in AID-RING1B line in response to auxin treatment, and that RING1B levels remained unchanged in auxin-treated TIR1-only control cells.

HaloTag-SUZ12 PRC1^CPM^ and RING1B-HaloTag PRC1^CPM^ lines were generated from PRC1^CPM^ ESCs using CRISPR-Cas9 guides specific for the N-terminus of *Suz12* and C-terminus of *Ring1b*, respectively, along with targeting constructs containing the HaloTag flanked by homology arms of at least 750 bp to introduce the HaloTag at both copies of each gene. After selection with puromycin, cells were labelled with 500 nM Halo-TMR, and cells with significantly higher fluorescence than the similarly labelled parental cell line were FACS-selected and plated at low density to allow picking of individual clones. Homozygous knock-in clones were identified by PCR screening, followed by western blot analysis to confirm homozygous tagging and that levels of protein expression were unchanged compared to the parental cell line.

### Calibrated ChIP-sequencing (cChIP-seq)

For RING1B, SUZ12, JARID2, PCL2, PCGF2, PCGF1 and PCGF6, cChIP-seq was performed as described previously (Fursova et al., 2019). Briefly, 5 x 10^7 mouse ESCs (untreated or following 72 hours OHT treatment) were mixed with 2 x 10^6 human HEK293T cells. Cells were resuspended in 10 ml phosphate buffered saline (PBS) and crosslinked at 25°C with 2 mM DSG (Thermo Scientific) for 45 minutes, and then with 1% formaldehyde (methanol-free, Thermo Scientific) for a further 15 minutes. Reactions were quenched with 125 mM glycine. Crosslinked cells were incubated in lysis buffer (50 mM HEPES pH 7.9, 140 mM NaCl, 1 mM EDTA, 10% glycerol, 0.5% NP40, 0.25% Triton-X100) for 10 minutes at 4°C. Released nuclei were washed (10 mM Tris-HCl pH 8, 200 mM NaCl, 1 mM EDTA, 0.5 mM EGTA) for 5 minutes at 4°C. Chromatin was then resuspended in 1 ml of sonication buffer (10 mM Tris-HCl pH 8, 100 mM NaCl, 1 mM EDTA, 0.5 mM EGTA, 0.1% Na deoxycholate, 0.5% N-lauroylsarcosine) and sonicated for 30 minutes using a BioRuptor Pico (Diagenode), shearing genomic DNA to an average size of 0.5 kb. Following sonication, TritonX-100 was added to a final concentration of 1%.

For ChIP, sonicated chromatin was diluted 10-fold in ChIP dilution buffer (1% Triton-X100, 1 mM EDTA, 20 mM Tris-HCl pH 8, 150 mM NaCl) and pre-cleared for 1 hour using Protein A agarose beads (Repligen) blocked with 1 mg/ml BSA and 1 mg/ml yeast tRNA. For each ChIP reaction, 1 ml of diluted and pre-cleared chromatin was incubated overnight with the appropriate antibody, anti-RING1B (CST, D22F2, 3 μl), anti-SUZ12 (CST, D39F6, 3 μl), anti-PCGF1 (in-house, 5 μl), anti-PCGF2 (Santa Cruz, sc-10744, 3 μl), anti-JARID2 (CST D6M9X, 3 μl), anti-PCL2 (GenWay GWB-FA7207, 2 μl), or anti-PCGF6 (3 μl). Antibody-bound chromatin was captured using blocked protein A agarose for 1 hour at 4°C and collected by centrifugation. ChIP washes were performed as described previously (Farcas et al., 2012). ChIP DNA was eluted in elution buffer (1% SDS, 0.1 M NaHCO3) and cross-links were reversed overnight at 65°C with 200 mM NaCl and 2 μl RNase A (Sigma). A matched input sample (10% of original ChIP reaction) was identically treated. The following day, ChIP samples and Inputs were incubated with Proteinase K (Sigma) for 1.5 hours at 56°C and purified using ChIP DNA Clean and Concentrator Kit (Zymo Research).

cChIP-seq libraries for both ChIP and Input samples were prepared using NEBNext Ultra DNA Library Prep Kit for Illumina, following manufacturer’s guidelines. Samples were indexed using NEBNext Multiplex Oligos. The average size and concentration of all libraries was analysed using the 2100 Bioanalyzer High Sensitivity DNA Kit (Agilent) followed by qPCR using SensiMix SYBR (Bioline, UK) and KAPA Illumina DNA standards (Roche). Libraries were sequenced as 40 bp paired-end reads on Illumina NextSeq 500 platform.

### Native cChIP-sequencing

Native cChIP-seq for H2AK119ub1, H3K27me3 and H3K4me3 was performed as described previously (Fursova et al., 2019). Briefly, 5 x 10^7 mouse ESCs (both untreated and following 72 hours OHT treatment) were mixed with 2 x 10^7 Drosophila SG4 cells in PBS. Mixed cells were pelleted and nuclei were released by resuspending in ice cold lysis buffer (10 mM Tris-HCl pH 8.0, 10 mM NaCl, 3 mM MgCl_2_, 0.1% NP40, 5 mM N-ethylmaleimide). Nuclei were then washed, and resuspended in 1 ml of MNase digestion buffer (10 mM Tris-HCl pH 8.0, 10 mM NaCl, 3 mM MgCl_2_, 0.1% NP40, 0.25 M sucrose, 3 mM CaCl_2_, 10 mM N-ethylmaleimide, 1x protease inhibitor cocktail (Roche)). Each sample was incubated with 200 units of MNase (Fermentas) at 37°C for 5 min, followed by the addition of 4 mM EDTA to halt MNase digestion. Following centrifugation at 1500 g for 5 min at 4°C, the supernatant (S1) was retained. The remaining pellet was incubated with 300 µl of nucleosome release buffer (10 mM Tris-HCl pH 7.5, 10 mM NaCl, 0.2 mM EDTA, 1x protease inhibitor cocktail (Roche), 10 mM N-ethylmaleimide) at 4°C for 1 h, passed five times through a 27G needle using a 1 ml syringe, and spun at 1500 g for 5 min at 4°C. The second supernatant (S2) was collected and combined with corresponding S1 sample from above. A small amount of S1/S2 DNA was purified and visualized on a 1.5% agarose gel to confirm digestion to mostly mono-nucleosomes.

For ChIP experiments, S1/S2 nucleosomes were diluted 10-fold in native ChIP incubation buffer (70 mM NaCl, 10 mM Tris pH 7.5, 2 mM MgCl2, 2 mM EDTA, 0.1% Triton, 1x protease inhibitor cocktail (Roche), 10 mM N-ethylmaleimide (NEM)), and 1 ml aliquots were made. Each ChIP reaction was then incubated overnight at 4°C with the appropriate antibody, 5 μl of anti-H2AK119ub1 (Cell Signaling Technology, D27C4), 5 μl of anti-H3K27me3 (in-house) or 3 μl anti-H3K4me3 (in-house) antibody. Antibody-bound nucleosomes were captured using protein A agarose (Repligen) beads, pre-blocked in native ChIP incubation buffer supplemented with 1 mg/ml BSA and 1 mg/ml yeast tRNA, for 1 hour at 4°C and collected by centrifugation. Immunoprecipitated material was washed four times with Native ChIP wash buffer (20 mM Tris pH 7.5, 2 mM EDTA, 125 mM NaCl, 0.1% Triton-X100) and once with Tris-EDTA buffer (10 mM Tris pH 8, 1 mM EDTA). ChIP DNA was eluted using 100 μl of elution buffer (1% SDS, 0.1 M NaHCO3), and then purified using ChIP DNA Clean and Concentrator Kit (Zymo Research). For each individual ChIP sample, DNA from a matched Input control (corresponding to 10% of original ChIP reaction) was also purified. Native cChIP-seq library preparation and sequencing was performed as described above for cChIP-seq.

### Calibrated RNA-sequencing (cRNA-seq)

For cRNA-seq, 1 x 10^7 mouse ESCs (both untreated and following 72 hours OHT treatment) were mixed with 4 x 10^6 Drosophila SG4 cells in 600 μl PBS. For RNA extraction, 400 μl of cells was used (corresponding to 6.7 x 10^6 mouse ESCs), and for DNA extraction the remaining 200 μl of cells was used (corresponding to 3.3 x 10^6 mouse ESCs). RNA extraction was performed using RNeasy mini kit columns (Qiagen) following manufacturer’s guidelines, and 10 μg was subjected to Turbo DNase (ThermoFisher) treatment to remove any contaminating DNA. Quality of RNA was assessed using 2100 Bioanalyzer RNA 6000 Pico kit (Agilent). Next, RNA samples were depleted of rRNA using the NEBNext rRNA Depletion kit (NEB). RNA-seq libraries were prepared using the NEBNext Ultra II Directional RNA Library Prep kit (NEB). To quantitate the consistency of spike-in cell mixing for each individual sample, a matched sample of cells was used to isolate genomic DNA using Quick-DNA miniprep kit (Zymo). Libraries from gDNA were prepared using NEBNext Ultra II FS kit (NEB) following manufacturer’s guidelines. RNA and DNA libraries were sequenced as 80 bp paired-end reads on the Illumina NextSeq 500 platform.

### Preparation of nuclear and histone extracts and immunoblotting

For nuclear extraction, ESCs were washed with PBS and then resuspended in 10 volumes of Buffer A (10 mM Hepes pH 7.9, 1.5 mM MgCl2, 10 mM KCl, 0.5 mM DTT, 0.5 mM PMSF and protease inhibitor cocktail (Roche)). After 10 min incubation on ice, cells were recovered by centrifugation at 1500 g for 5 min and resuspended in 3 volumes of Buffer A supplemented with 0.1% NP-40. The released nuclei were pelleted by centrifugation at 1500 g for 5 min, followed by resuspension in 1 volume of Buffer C (5 mM Hepes (pH 7.9), 26 % glycerol, 1.5 mM MgCl 2, 0.2 mM EDTA, protease inhibitor cocktail (Roche) and 0.5 mM DTT) supplemented with 400 mM NaCl. The extraction was allowed to proceed on ice for 1 hour with occasional agitation, then the nuclei were pelleted by centrifugation at 16,000 g for 20 min at 4°C. The supernatant was taken as the nuclear extract.

For histone extraction, ESCs were washed with RSB supplemented with 20 mM NEM, incubated on ice for 10 min in RSB with 0.5% NP-40 and 20 mM NEM, pelleted by centrifugation at 800 g for 5 min and incubated in 2.5 mM MgCl2, 0.4 M HCl and 20 mM NEM on ice for 30 min. After that, cells were pelleted by centrifugation at 16,000 g at 4°C for 20 min, the supernatant recovered and precipitated on ice with 25% TCA for 30 min, followed by centrifugation at 16,000 g for 15 min at 4°C to recover histones. Following two acetone washes, the histones were resuspended in 150 μl 1xSDS loading buffer and boiled at 95°C for 5 min. Finally, any insoluble precipitate was pelleted by centrifugation at 16,000 g for 15 min at 4°C and the soluble fraction retained as the histone extract. Histone concentrations across samples were compared by Coomassie Blue staining following SDS-PAGE. Semi-quantitative western blot analysis of histone extracts was performed using LI-COR IRDye® secondary antibodies and imaging was done using the LI-COR Odyssey Fc system. To measure the changes in bulk H2AK119ub1 levels, the relative signal of H2AK119ub1 to H3 or H4 histones was quantified.

### Co-immunoprecipitation

For co-immunoprecipitation reactions, 400 μg of nuclear extract from wild-type or RING1B^I53A/D56K^ ESCs was added to BC150 buffer (150 mM KCl, 10% glycerol, 50 mM HEPES (pH 7.9), 0.5 mM EDTA, 0.5 mM DTT) with 1x protease inhibitor cocktail (Roche) to a total volume of 550 μl. A 50 μl Input sample was retained, and 5 μg of mouse monoclonal anti-RING1B antibody (Atsuta et al., 2001) was added to the remaining 500 μl of sample. Immunoprecipitation reactions were then incubated overnight at 4°C. Immunoprecipitated material was collected with Protein A agarose beads and washed four times in 1 ml of BC150 buffer. Following the final wash step, beads were directly resuspended in 100 μl of 1x SDS loading buffer (2% SDS, 0.1 M Tris pH 6.8, 0.1 M DTT, 10% glycerol, 0.1% bromophenol blue) and placed at 95°C for 5 mins. 1x SDS loading buffer was similarly added to Input samples which were also incubated at 95°C for 5 mins, prior to SDS-PAGE and western blot analysis.

### Polycomb body imaging

To image Polycomb bodies in live cells, HaloTag-SUZ12;PRC1^CPM^ or RING1B-HaloTag;PRC1^CPM^ cells were plated on gelatinised 35 mm petri dish, 14 mm Microwell 1.5 coverglass dishes (MatTek, #P35G-1.5-14-C) at least 5 hours before imaging. Prior to imaging, cells were labelled with 500 nm JF_549_ (Grimm et al., 2017) for 15 min at 37°C, followed by 3 washes, changing medium to Fluorobrite DMEM (Thermo Fisher Scientific) supplemented as described for general ESC culture above. Cells were incubated for a further 30 min in supplemented Fluorobrite DMEM with 10 µg/mL Hoechst 33258 (Thermo Fisher Scientific) at 37°C and washed once more before imaging. Cells were imaged on an IX81 Olympus microscope connected to a Spinning Disk Confocal system (UltraView VoX PerkinElmer) using an EMCCD camera (ImagEM, Hamamatsu Photonics) in a 37°C heated, humidified, CO_2_-controlled chamber. Z-stacks were acquired using a 100x PlanApo NA 1.40 oil-immersion objective heated to 37°C, using Volocity software (PerkinElmer). HaloTag-JF_549_ was imaged with a 561 nm laser at 1.25 sec exposure at 15% laser power, while Hoechst was imaged with a 405 nm laser at 250 ms exposure at 20% laser power. Z-stacks were acquired at 150 nm intervals.

### Capture-C library preparation

CaptureC libraries were prepared as described previously (Hughes et al., 2014). Briefly, 10^6^ mouse ESCs were trypsinized, collected in 50 ml falcon tubes in 9.3 ml media and crosslinked with 1.25 ml 16% formaldehyde for 10 minutes at room temperature. Cells were quenched with 1.5 M glycine, washed with PBS and lysed for 20 minutes at 4°C lysis buffer (10mM Tris pH 8, 10 mM NaCl, 0.2% NP-40, supplemented with complete proteinase inhibitors) prior to snap freezing in 1 ml lysis buffer at −80°C. Lysates were then thawed on ice, pelleted and resuspended in 650 µl 1x *Dpn*II buffer (NEB). Three 1.5ml tubes with 200µl lysate each were treated in parallel with SDS (0.28% final concentration, 1hr, 37°C, interval shaking 500 rpm, 30 sec on/30 sec off), quenched with trypsin (1.67% final concentration, 1hr, 37°C, interval shaking 500rpm, 30 sec on/30 sec off) and subjected to a 24 hour digestion with 3×10µl DpnII (homemade, 37°C, interval shaking 500rpm, 30 sec on/30 sec off). Each chromatin aliquot was independently ligated with 8 µl T4 Ligase (240 U) in a volume of 1440 µl (20 hours, 16°C). Following this, the nuclei containing ligated chromatin were pelleted to remove any non-nuclear chromatin, reverse-crosslinked and the ligated DNA was phenol-chloroform purified. The sample was resuspended in 300 µl water and sonicated 13x (Pico Bioruptor, 30 sec on, 30 sec off) or until a fragment size of approximately 200 bp was reached. Fragments were size-selected using AmpureX beads (Beckman Coulter: A63881, ratios: 0.85x / 0.4x). 2x 1-5 µg of DNA were adaptor ligated and indexed using the NEBNext DNA library Prep Reagent Set (New England Biolabs: E6040S/L) and NEBNext Multiplex Oligos for Illumina Primer sets 1 (New England: E7335S/L) and 2 (New England: E7500S/L). The libraries were amplified 7x using Herculase II Fusion Polymerase kit (Agilent: 600677).

### Capture-C hybridization and sequencing

5’ biotinylated probes were designed using the online tool by the Hughes lab (CapSequm) to be 70-120bp long and two probes for each promoter of interest. The probes were pooled at 2.9 nM each. Samples were captured twice and hybridizations were carried out for 72h and for 24h for the first and the second captures, respectively. To even out capture differences between tubes, libraries were pooled prior to hybridization at 1.5 µg each. Hybridization was carried out using Nimblegen SeqCap (Roche, Nimblegen SeqCap EZ HE-oligo kit A, Nimblegen SeqCap EZ HE-oligo kit B, Nimblegen SeqCap EZ Accessory kit v2, Nimblegen SeqCap EZ Hybridisation and wash kit) following manufacturer’s instructions for 72 hours followed by a 24 hours hybridization (double Capture). The captured library molarity was quantified by qPCR using SensiMix SYBR (Bioline, UK) and KAPA Illumina DNA standards (Roche) and sequenced on Illumina NextSeq 500 platform as 80 bp paired-end reads for three biological replicates.

### Massive parallel sequencing, data processing and normalisation

For calibrated ChIP-seq, paired-end reads were aligned to the concatenated mouse and spike-in genome sequences (mm10+dm6 for native cChIP-seq, and mm10+hg19 for cross-linked cChIP-seq) using Bowtie 2 (Langmead and Salzberg, 2012) with the “--no-mixed” and “--no-discordant” options. Only uniquely mapped reads after removal of PCR duplicates with Sambamba (Tarasov et al., 2015) were used for downstream analysis.

For cRNA-seq, first, paired-end reads were aligned using Bowtie 2 (with “--very-fast”, “--no-mixed” and “--no-discordant” options) against the concatenated mm10 and dm6 rDNA genomic sequence (GenBank: BK000964.3 and M21017.1) and reads mapping to rDNA were discarded. All unmapped reads were then aligned against the concatenated mm10 and dm6 genome sequences using STAR (Dobin et al., 2013). Finally, reads that failed to map using STAR were aligned against the mm10+dm6 concatenated genome using Bowtie 2 (with “--sensitive-local”, “--no-mixed” and “--no-discordant” options) to improve mapping of introns. Uniquely aligned reads from the last two steps were combined for further analysis. PCR duplicates were removed using Sambamba (Tarasov et al., 2015). For the corresponding gDNA-seq experiments, paired-end read alignment and processing was carried out as described above for cChIP-seq.

For Capture-C, paired-end reads were aligned and filtered for HiC artefacts using HiCUP (Wingett et al., 2015) and Bowtie2 (Langmead and Salzberg, 2012) with fragment filter set to 100-800bp. A list of all Next-Generation sequencing experiments carried out in this study and the number of uniquely aligned reads in each experiment can be found in Table S2.

For visualisation of cChIP-seq and cRNA-seq data and annotation of genomic regions with read counts, uniquely aligned mouse reads were normalised using dm6 or hg19 spike-in as described previously (Fursova et al., 2019). Briefly, mm10 reads were randomly subsampled based on the total number of spike-in (dm6 or hg19) reads in each sample. To account for any minor variations in spike-in cell mixing between replicates, we used the ratio of spike-in/mouse total read counts in the corresponding Input/gDNA-seq samples to correct the subsampling factors. After normalisation, read coverages for individual biological replicates were compared across RING1B peaks for cChIP-seq or gene bodies for cRNA-seq using multiBamSummary and plotCorrelation from deepTools (Ramirez et al., 2014). For each experimental condition, biological replicates correlated well (Pearson correlation coefficient > 0.9, see Table S3) and were merged for downstream analysis. Genome coverage tracks were generated using the pileup function from MACS2 (Zhang et al., 2008) for cChIP-seq and genomeCoverageBed from BEDTools (Quinlan, 2014) for cRNA-seq and visualised using the UCSC genome browser (Kent et al., 2002). BigwigCompare from deeptools (v3.1.1) (Ramirez et al., 2014) was used to make differential genome coverage tracks.

### Peak calling

To identify genomic regions bound by RING1B, SUZ12 and PCGF2, we carried out peak calling using MACS2 (“BAMPE” and “--broad” options specified), with corresponding Input samples used as a control. For RING1B and SUZ12, peaks were called using merged biological replicates from untreated PRC1^CPM^ and PRC1^CKO^ cells for RING1B and SUZ12 cChIP-seq respectively, and only peaks identified in both cell lines were selected. For PCGF2, individual replicates from untreated PRC1^CPM^ ESC were used, and only peaks identified in all biological replicates were taken forward. For all peak sets, peaks overlapping with a custom-build set of blacklisted genomic regions were discarded to remove sequencing artefacts. For RING1B-bound regions, RING1B cChIP-seq in PRC1^CKO^ ESCs was used to filter out peaks which showed no significant loss of RING1B signal following tamoxifen treatment (p-adj < 0.05 and > 2-fold). In total, we were able to identify 18643 RING1B peaks, 7438 SUZ12 peaks and 3680 PCGF2 peaks. Using the overlap between these peak sets, we defined Polycomb (PcG)-occupied regions as RING1B peaks that overlap SUZ12 peaks (n = 7074), and PCGF2 target sites as RING1B peaks overlapping PCGF2 peaks (n = 3568). To characterise low-level genomic blanket of H2AK119ub1, we have generated a set of 100 kb windows spanning the genome (n = 27,348) using makewindows function from BEDtools (v2.17.0).

### Read count quantitation and analysis

For cChIP-seq, computeMatrix and plotProfile/plotHeatmap from deeptools were used to perform metaplot and heatmap analysis of read density at regions of interest. For each cChIP-seq dataset and each cell line, read density at the peak centre in untreated cells was set to 1. For chromosome-wide density plots, read coverage in 250 kb bins was calculated using a custom R script utilising GenomicRanges, GenomicAlignments and Rsamtools Bioconductor packages (Huber et al., 2015) and visualised using ggplot2. For cChIP-seq, target regions of interest were annotated with read counts using multiBamSummary from deeptools (“--outRawCounts”). For comparative box plot analysis, read counts from merged spike-in normalised replicates were used, while for differential enrichment analysis, read counts from individual biological replicates prior to spike-in normalisation were obtained. For differential gene expression analysis, we used a custom Perl script utilising SAMtools to obtain read counts for a custom-built non-redundant mm10 gene set (n = 20633), derived from mm10 refGene genes by removing very short genes with poor sequence mappability and highly similar transcripts.

Normalised read counts and log2 fold changes for different genomic intervals were visualised using custom R scripts and ggplot2. For box plot analysis of cChIP-seq signal for different factors before and after treatment, read counts were normalised to the genomic region size (in kb) and median value of cChIP-seq signal in untreated cells, log2 of which was set to 1. For box plots comparing H2AK119ub1 enrichment at RING1B-bound sites and 100 kb genomic windows, read density is shown relative to the signal at RING1B-bound sites in untreated cells. For box plots, boxes show interquartile range (IQR) and whiskers extend by 1.5xIQR. To study the relationship between the levels of different factors/histone modifications and gene expression at RING1B-bound genomic regions, the RING1B target sites that overlapped with gene promoters were divided into percentiles based on the expression level of the associated gene in untreated PRC1^CPM^ cells. For each percentile, mean read density normalised to the genomic region size (in kb) was plotted relative to the mean read density in the fiftieth percentile, together with loess regression trendlines. ggcor function from the GGally (v1.4.0) R package was used to generate a correlation matrix for the association of RING1B occupancy before and after tamoxifen treatment in PRC1^CPM^ cells with a set of chromatin features and occupancy of other Polycomb factors. All correlation analyses in the paper used Pearson correlation coefficient to measure the strength of the association between the variables and were visualised using scatterplots coloured by density with stat_density2d. Linear regression lines were plotted using stat_poly_eq function from the ggpmisc (v0.3.1) R package, together with the model’s R^2^coefficient of determination. Student’s t-test and Wilcoxon signed-rank statistical tests were also performed in R with samples considered to be independent and two-sided alternative hypothesis, unless otherwise specified.

For CaptureC, read counts and interaction scores (significance of interactions) for the captured gene promoters were obtained using the Bioconductor package CHiCAGO (Cairns et al., 2016). For visualisation of CaptureC data, weighted read counts from CHiCAGO data files for merged biological replicates were normalized to the total number of reads aligning to the captured gene promoters and further to the number of promoters in the respective CaptureC experiment. Bigwig files were generated from these normalized read counts. For comparative boxplot analysis, interactions called by CHiCAGO (score >= 5) across all samples were aggregated and interactions with a distance of less than 10 DpnII fragments were merged to a single interaction peak. For each interaction peak, we then quantified mean normalized read count and CHiCAGO scores of all overlapping *Dpn*II fragments.

### Differential cChIP-seq enrichment and gene expression analysis

To identify significant changes in cChIP-seq and cRNA-seq, we used a custom R script that incorporates spike-in calibration into DESeq2 analysis (Love et al., 2014). In order to do this, we used read counts from the spike-in genome at a control set of intervals to calculate DESeq2 size factors for normalisation of raw mm10 read counts (as has been previously described in (Fursova et al., 2019; Taruttis et al., 2017). Unique dm6 genes from refGene were used for spike-in normalisation of cRNA-seq, while 10 kb (for hg19 spike-in) or 1 kb (for dm6 spike-in) windows spanning the genome were used for differential enrichment analysis of cChIP-seq. Prior to quantification, spike-in reads were pre-normalised using the spike-in/mouse read ratio derived from the corresponding Input or genomic DNA-seq sample in order to account for minor variations in mixing of mouse and spike-in cells. For a change to be called significant, we applied a threshold of p-adj < 0.05 and fold change > 1.5, unless otherwise specified. Log2-fold change values were visualised using R and ggplot2 with MA plots and violin plots. For MA plots, density of the data points across y-axis is shown to reflect the general direction of gene expression changes.

### Annotation of Polycomb target genes

Mouse genes in a custom non-redundant set (n = 20633) that was used for differential gene expression analysis were classified into three groups based on the overlap of their gene promoters with non-methylated CpG islands (NMI), as well as RING1B- and SUZ12-bound sites. NMIs (n = 27047) were identified using MACS2 peak calling from BioCAP-seq data (Long et al., 2013) as regions which are enriched with non-methylated CpG-rich DNA. All genes with promoters (TSS ± 2500 bp) not overlapping with NMIs were referred to as non-NMI genes (n = 5326). Genes that contained NMIs at their promoters were further sub-divided into PcG-occupied genes (n = 6283), if their promoters also overlapped with both RING1B and SUZ12-bound sites defined in this study, and non-PcG-occupied genes (n = 9024), if they did not. For genes with several described transcripts and promoters, a complete mm10 refGene gene set was used for classification, with the overlap of at least one promoter with the feature of interest being required to assign a gene into PcG/non-PcG occupied categories. Finally, we refer to a subset of PcG-occupied genes which showed a statistically significant increase in gene expression following removal of PRC1 in PRC1^CKO^ cells, as PRC1-repressed genes (n = 2482).

In order to identify genes that were differentially affected by the removal of PRC1 or specifically PRC1 catalytic activity, we used two complementary approaches. First, we isolated genes that were expressed at significantly lower levels (p-adj < 0.05 and fold change > 1.5) in PRC1^CPM^ as compared to PRC1^CKO^ cells following tamoxifen treatment. In addition, we also identified genes, for which the magnitude of expression changes following tamoxifen treatment was significantly smaller in PRC1^CPM^ cells than in PRC1^CKO^ cells, using the DESeq2 design that included the interaction term between the cell line and treatment factors. Combination of these two approaches has yielded a total of 241 PRC1-repressed genes that were derepressed to a smaller extent in PRC1^CPM^ cells as compared to PRC1^CKO^ cells following tamoxifen treatment.

### Analysis of live-cell imaging of Polycomb bodies

To segment Polycomb bodies in individual live cells for analysis, nuclei were first manually segmented based on Hoechst fluorescence using TANGO in ImageJ (Ollion et al., 2013). 561 nm channels of z-stacks were deconvolved using Olympus cellSens software (constrained iterative deconvolution, 5 cycles). Deconvolved 561 nm z-stacks were masked using outputs from TANGO, and individual Polycomb bodies identified using a custom script. Briefly, segmented nuclei were background subtracted using a 4 px rolling ball and a mask of Polycomb bodies generated using Otsu thresholding. 3D Objects Counter in ImageJ was used to quantify the properties of the masked Polycomb bodies, and its outputs were processed and analysed using R.

### Data and software availability

The high-throughput data reported in this study have been deposited in GEO under the accession number GSEXXXXXX. Published data used in this study include BioCAP-seq (GSE43512 (Long et al., 2013)); MAX ChIP-seq (GSE48175; (Krepelova et al., 2014); H2AK119ub1, H3K27me3 and RING1B cChIP-seq together with corresponding Inputs in PRC1^CKO^ ESCs (GSE119618; (Fursova et al., 2019)); cRNA-seq in *Pcgf4^-/-^4;Pcgf2^fl/fl^*(GSE119619; (Fursova et al., 2019)); CaptureC in RING1B^deg^ and control cell lines (ArrayExpress E-XXXX-XXXX (Rhodes et al., 2019)). All R and Perl scripts used for data analysis in this study are available upon request.

